# In silico analysis of *IDH3A* gene revealed Novel mutations associated with Retinitis Pigmentosa

**DOI:** 10.1101/554196

**Authors:** Thwayba A. Mahmoud, Abdelrahman H. Abdelmoneim, Naseem S. Murshed, Zainab O. Mohammed, Dina T. Ahmed, Fatima A. Altyeb, Nuha A. Mahmoud, Mayada A. Mohammed, Fatima A. Arayah, Wafaa I. Mohammed, Omnia S. Abayazed, Amna S. Akasha, Mujahed I. Mustafa, Mohamed A. Hassan

## Abstract

**Background:** Retinitis Pigmentosa (RP) refers to a group of inherited disorders characterized by the death of photoreceptor cells leading to blindness. The aim of this study is to identify the pathogenic SNPs in the IDH3A gene and their effect on the structure and function of the protein.

**Method:** we used different bioinformatics tools to predict the effect of each SNP on the structure and function of the protein.

**Result:** 20 deleterious SNPs out of 178 were found to have a damaging effect on the protein structure and function.

**Conclusion:** this is the first in silico analysis of IDH3A gene and 20 novel mutations were found using different bioinformatics tools, and they could be used as diagnostic markers for Retinitis Pigmentosa.

## 1. Introduction

Retinitis Pigmentosa (RP) refers to a group of inherited disorders characterized by the death of photoreceptor cells (rods and cons) leading to blindness (1–12) It affects about 2.5 million people worldwide (5) and symptoms usually starts with night blindness followed by vision loss, progressive restriction of the peripheral visual field and abnormalities in the electroretinogram (1–4, 13). RP is classified as nonsyndromic (70%-80%) (14, 15), syndromic or systemic (4, 16). The most frequent form of syndromic RP is Usher syndrome (14, 17) which involves neurosensory hearing loss, and Bardet-Biedl syndrome (18, 19) which involves RP, obesity, renal abnormalities and mental retardation.

RP has three patterns of inheritance; autosomal dominant (adRP), autosomal recessive (arRP) and X-linked pattern (14, 20–22) and the most reported genes for each case are *RHO* gene, *USH2A* and *RPGR* respectively (4, 21–24) yet the gene we are working on, Isocitrate Dehydrogease 3, (*IDH3A*) is a novel gene identified as the cause of typical arRP (25) located in the human chromosome 15q25.1 (26). The *IDH3A* function is to catalyze the oxidative decarboxylation of isocitrate (27) which is the key rate-limiting step of the tricarboxylic acid cycle. It also plays a central role in the aerobic energy production and cellular respiration thus mutations in this gene will affect the nervous system (28, 29) and may cause cancer (30, 31).

Although the RP is the most common inherited disease of the retina (14, 32–36) there is still no effective therapeutic strategy and the exact pathogenesis and etiology of the disease is not clear (3, 5, 32, 37).

The aim of this study is to identify the pathogenic SNPs in the *IDH3A* gene and their effect on the structure and function of the protein, which might help in the overall understanding of the pathogenesis of the disease and could be used as diagnostic markers. This is the first in silico analysis in the coding region of *IDH3A* gene to prioritize SNPs for further genetic mapping studies. Utilization of in analysis softwares expedites the process of identifying the deleterious SNPs with no cost, and also facilitates future genetic studies.(38)

## 2. Method

### 2.1 Data Mining

The data concerning the *IDH3A* human gene was procured from the National Center for Biotechnology Information (NCBI)(http://www.ncbi.nlm.nih.gov/) (39). The SNPs information of the *IDH3A* gene was collected from the NCBI dbSNP database (http://www.ncbi.nlm.nih.gov/snp/) (40) and the IDH3A protein sequence was obtained from UniProt database (https://www.uniprot.org/) (41).

### 2.2 SIFT

SIFT algorithm predicts damaging and tolerated (non-damaging) substitutions based on sequence homology and physical properties of sequence submitted. It also predicts the functional consequences of amino acid substitutions caused due to nsSNPs. SIFT is based on the premise that protein evolution is correlated with protein function. It uses multiple alignment information to predict tolerated and deleterious substitutions for every position of the sequence of interest. (42, 43)

SIFT score ranges from 0 to 1. The amino acid substitution is predicted to be damaging if the score is ≤ 0.05, and tolerated if the score is > 0.05. (https://sift.bii.a-star.edu.sg/)

### 2.2 PolyPhen-2

PolyPhen-2 (polymorphism phenotyping v2), available as software and via a Web server, predicts the possible impact of amino acid substitutions on the stability and function of human proteins using structural and comparative evolutionary considerations. PolyPhen-2 uses eight sequence-based and structure-based predictive features which were selected automatically by an interactive greedy algorithm. Majority of these features involves comparison of a property of the wild-type residue and the corresponding property of the mutant.

PolyPhen-2 uses multiple sequence alignment by selecting the homologues sequences for the analysis using a clustering algorithm and then constructs and refines their multiple alignments. The functional significance of an amino acid replacement is predicted from its individual features by Naïve Bayes classifier. (44, 45)

Prediction outcomes are benign, possibly damaging or probably damaging according to the PSIC value which ranges from 0 to 1. Values closer to zero are considered benign while values closer to one are considered probably damaging. PolyPhen-2 is available at (http://genetics.bwh.harvard.edu/pph2/).

### 2.3 PROVEAN

Provean predicts the functional effect of protein sequence variations, including single amino acid substitutions and small insertions and deletions. The prediction is based on the change, caused by a given variation, in the similarity of a query sequence to a set of its related protein sequences. For this prediction the algorithm is required to compute a semi global pair-wise sequence alignment score between the query sequence and each of the related sequences (46, 47). Provean is available at this website (http://provean.jcvi.org/index.php).

### 2.4 PhD-SNP

A predictor of human deleterious SNPs based on an online Support Vector Machine (SVM) classifier explicitly designed for human dataset correlated with disease inducing mutations. The SVM classifies mutations into disease related (desired output set to 0) and neutral polymorphism, based on protein sequence and profile information. The reliability index value (RI) is evaluated and the probability of the amino acid being deleterious is obtained. A probability> 0.5 is considered disease associated whereas ≤0.5 is considered neutral. PhD-SNP is available at this website (http://snps.biofold.org/phd-snp/phd-snp.html).

### 2.5 SNP&GO

SNP&GO is a method for the prediction of deleterious SNPs using protein functional annotation. The server is based on SVM classifier that discriminates between disease related and neutral SNPs. It has two components, one is sequence based and the other is structure based. The output gives the mutated residue, the prediction (either disease or neutral), RI (probability of disease related class) and the information about the prediction method. If the probability is > 0.5 then the variation is disease associated. SNP&GO is available at this website (http://snps.biofold.org/snps-and-go/index.html) (48, 49).

### 2.6 P-Mut

P-Mut uses a robust methodology to predict disease-associated mutations. It allows fast and accurate prediction based on the use of neural networks (NNs) trained with a large database of neutral mutations and pathological mutations of mutational hot spots which are obtained by alanine scanning, massive mutations and genetically accessible mutations. The final output is displayed as a pathogenecity index ranging from 0 to 1 and a confidence index ranging from 0 to 9(50). P-Mut server is available at this website (http://mmb.irbbarcelona.org/PMut/).

### 2.7 I-Mutant 3.0

I-Mutant is an SVM-based tool for the automatic prediction of protein stability changes upon single point mutations. The predictions are performed starting either from the protein structure or, more importantly, from the protein sequence. I-Mutant correctly predicts 80% or 77% of the dataset, depending on the usage of structural or sequence information, respectively. The server was also trained to predict the value of the free energy stability change upon single point mutation, starting from the protein structure or sequence (51). (http://gpcr2.biocomp.unibo.it/cgi/predictors/I-Mutant3.0/I-Mutant3.0.cgi)

### 2.8 MUpro

MUpro is a SVM-based tool for the prediction of protein stability changes upon nsSNPs. The value of the energy change is predicted, and a confidence score between −1 and 1 for measuring the confidence of the prediction is calculated. The accuracy for SVM using sequence information is 84.2%. MUpro is available at (http://mupro.proteomics.ics.uci.edu/).

### 2.9 Chimera 1.8

Chimera is a highly extensible program of interactive visualization and analysis of molecular structures and related data, including density maps, supramolecular assemblies, sequence alignments, docking results, trajectories and conformational ensembles. High quality images and animations can be generated. Chimera is segmented into a core that provides basic services and visualization, and extensions that provides higher level functionalities (52). The protein structure was obtained from RaptorX server (http://raptorx.uchicago.edu/). Chimera is available at (http://www.cgl.ucsf.edu/chimera/).

### 2.10 GeneMania

GeneMania is a flexible, user-friendly web interface for generating hypothesis about gene function, analyzing gene lists and prioritizing genes for functional assays. Given a query gene list, GeneMania finds functionally similar genes using a wealth of genomics and proteomics data. Another use of GeneMania is gene function prediction by finding the genes that are likely to share functions with the query based on their interactions with it (53–55). (https://genemania.org/).

### 2.11 Project HOPE

Project HOPE is next-generation web application for automatic mutant analysis and it explains the molecular origin of a disease related phenotype caused by mutations in human proteins. HOPE collects information from data sources such as the protein’s 3D structure and the UniProt database of well-annotated protein sequences. A decision scheme is used to process these data and to predict the effect of the mutation on the 3D structure and the function of the protein. Then a report is produced which explains and illustrate the effect of the mutations (56). (http://www.cmbi.ru.nl/hope/method/)

### 2.12 BioEdit

BioEdit is a user-friendly sequence alignment editor which enables the user to manually edit a multiple sequence alignment in order to obtain a more reasonable or expected alignment. The sequences are entered in a FastA format and the output file is provided in a suitable user-designated format. BioEdit is available for download at (http://www.mbio.ncsu.edu/bioedit/bioedit.html).

## 3. Result

**Table (1):** Deleterious (damaging) nsSNPs predicted by various softwares:

| Amino acid change | SIFT |  | Polyphen-2 |  | PROVEAN |  |
| --- | --- | --- | --- | --- | --- | --- |
|  | prediction | score | prediction | score | prediction | score |
| G39V | affect | 0 | probably damaging | 1 | Deleterious | -8.482 |
| I46S | affect | 0 | probably damaging | 1 | Deleterious | -5.655 |
| V50F | affect | 0 | probably damaging | 1 | Deleterious | -4.679 |
| G97C | affect | 0 | probably damaging | 1 | Deleterious | -8.624 |
| A122T | affect | 0 | probably damaging | 1 | Deleterious | -3.904 |
| A122S | affect | 0 | probably damaging | 1 | Deleterious | -2.908 |
| A122E | affect | 0 | probably damaging | 1 | Deleterious | -4.887 |
| T172I | affect | 0 | probably damaging | 1 | Deleterious | -5.824 |
| R178C | affect | 0 | probably damaging | 1 | Deleterious | -7.443 |
| R178H | affect | 0 | probably damaging | 1 | Deleterious | -4.459 |
| F184L | affect | 0 | probably damaging | 1 | Deleterious | -5.924 |
| Y186C | affect | 0 | probably damaging | 1 | Deleterious | -8.131 |
| V198G | affect | 0 | probably damaging | 1 | Deleterious | -6.794 |
| L210P | affect | 0 | probably damaging | 1 | Deleterious | -6.487 |
| P243R | affect | 0 | probably damaging | 1 | Deleterious | -8.773 |
| G268R | affect | 0 | probably damaging | 1 | Deleterious | -7.8 |
| G271S | affect | 0 | probably damaging | 1 | Deleterious | -5.827 |
| A302V | affect | 0 | Probably Damaging | 1 | Deleterious | -3.887 |
| P304H | affect | 0 | probably damaging | 1 | Deleterious | -8.715 |
| I327T | Affect | 0 | probably damaging | 1 | Deleterious | -4.766 |

**Table (2):** Disease-associated nsSNPs predicted by various softwares:

| sub | PhD-SNP<br>prediction | RI | score | SNP&GO<br>prediction | RI | score | P-Mut<br>prediction | score |
| --- | --- | --- | --- | --- | --- | --- | --- | --- |
| G39V | Disease | 8 | 0.914 | Disease | 7 | 0.826 | Disease | 0.93<br>(94%) |
| I46S | Disease | 7 | 0.864 | Disease | 1 | 0.552 | Disease | 0.93<br>(94%) |
| V50F | Disease | 7 | 0.829 | Disease | 7 | 0.854 | Disease | 0.93<br>(94%) |
| G97C | Disease | 8 | 0.898 | Disease | 7 | 0.835 | Disease | 0.93<br>(94%) |
| A122T | Disease | 4 | 0.706 | Disease | 4 | 0.679 | Disease | 0.70<br>(86%) |
| A122S | Disease | 5 | 0.757 | Disease | 4 | 0.686 | Disease | 0.75<br>(87%) |
| A122E | Disease | 7 | 0.868 | Disease | 6 | 0.813 | Disease | 0.93<br>(94%) |
| T172I | Disease | 7 | 0.841 | Disease | 5 | 0.74 | Disease | 0.93<br>(94%) |
| R178C | Disease | 8 | 0.914 | Disease | 6 | 0.788 | Disease | 0.93<br>(94%) |
| R178H | Disease | 7 | 0.857 | Disease | 4 | 0.704 | Disease | 0.88<br>(92%) |
| Y186C | Disease | 7 | 0.836 | Disease | 4 | 0.71 | Disease | 0.93<br>(94%) |
| V198G | Disease | 7 | 0.827 | Disease | 4 | 0.717 | Disease | 0.87<br>(91%) |
| L210P | Disease | 9 | 0.933 | Disease | 8 | 0.878 | Disease | 0.93<br>(94%) |
| P243R | Disease | 8 | 0.882 | Disease | 5 | 0.751 | Disease | 0.93<br>(94%) |
| G268R | Disease | 8 | 0.913 | Disease | 7 | 0.854 | Disease | 0.93<br>(94%) |
| G271S | Disease | 8 | 0.922 | Disease | 7 | 0.86 | Disease | 0.92<br>(94%) |
| P304H | Disease | 8 | 0.876 | Disease | 5 | 0.74 | Disease | 0.93<br>(94%) |
| I327T | Disease | 6 | 0.792 | Disease | 5 | 0.769 | Disease | 0.93<br>(94%) |

**Table (3):** The effect of the deleterious SNPs on the stability predicted by 2 softwares:

| Amino acid change | SVM2 Prediction Effect | RI | DDG Value Prediction | Mupro prediction | Mupro score |
| --- | --- | --- | --- | --- | --- |
| G39V | Decrease | 5 | -0.26 | Decrease | -0.95687971 |
| I46S | Decrease | 9 | -1.97 | Decrease | -1.2674836 |
| V50F | Decrease | 7 | -0.99 | Decrease | -1.4400617 |
| G97C | Decrease | 7 | -0.90 | Decrease | -0.37395268 |
| A122T | Decrease | 7 | -0.87 | Decrease | -0.91965426 |
| A122S | Decrease | 9 | -0.94 | Decrease | -0.75236358 |
| A122E | Decrease | 3 | -0.62 | Decrease | -0.46426725 |
| T172I | Decrease | 4 | -0.33 | Decrease | -0.15840056 |
| R178C | Decrease | 0 | -0.61 | Decrease | -0.63648418 |
| R178H | Decrease | 6 | -1.02 | Decrease | -1.1281043 |
| Y186C | Decrease | 0 | -1.13 | Decrease | -0.72393174 |
| V198G | Decrease | 9 | -2.09 | Decrease | -2.7354434 |
| L210P | Decrease | 8 | -2.08 | Decrease | -2.4575871 |
| P243R | Decrease | 4 | -0.87 | Decrease | -0.37063631 |
| G268R | Decrease | 3 | -0.48 | Decrease | -0.76962057 |
| G271S | Decrease | 3 | -0.97 | Decrease | -0.22864936 |
| P304H | Decrease | 7 | -1.35 | Decrease | -0.72552786 |
| I327T | Decrease | 8 | -1.67 | Decrease | -1.3879025 |

**Table (4):** *IDH3A*gene function and its appearance in network and genome:

| Function | FDR | Genes in network | Genes in genome |
| --- | --- | --- | --- |
| tricarboxylic acid cycle | 1.20E-12 | 7 | 21 |
| aerobic respiration | 2.19E-11 | 7 | 33 |
| energy derivation by oxidation of organic compounds | 4.26E-08 | 9 | 285 |
| cellular respiration | 4.30E-07 | 7 | 140 |
| mitochondrial matrix | 5.43E-07 | 8 | 257 |
| tricarboxylic acid metabolic process | 1.05E-06 | 4 | 11 |
| oxidoreductase activity, acting on the CH-OH group of donors, NAD or NADP as acceptor | 0.002837 | 4 | 73 |
| proteasome accessory complex | 0.002932 | 3 | 20 |
| oxidoreductase activity, acting on CH-OH group of donors | 0.003524 | 4 | 82 |
| regulation of cellular ketone metabolic process | 0.019321 | 4 | 129 |
| proteasome complex | 0.038133 | 3 | 51 |
| regulation of cellular amino acid metabolic process | 0.04157 | 3 | 57 |
| DNA damage response, signal transduction by p53 class mediator resulting in cell cycle arrest | 0.04157 | 3 | 67 |
| signal transduction involved in mitotic DNA damage checkpoint | 0.04157 | 3 | 68 |
| cell cycle arrest | 0.04157 | 4 | 191 |
| signal transduction involved in DNA integrity checkpoint | 0.04157 | 3 | 68 |
| negative regulation of ubiquitin-protein ligase activity involved in mitotic cell cycle | 0.04157 | 3 | 67 |
| signal transduction involved in mitotic G1 DNA damage checkpoint | 0.04157 | 3 | 67 |
| signal transduction involved in mitotic DNA integrity checkpoint | 0.04157 | 3 | 68 |
| regulation of cellular amine metabolic process | 0.04157 | 3 | 67 |
| cellular ketone metabolic process | 0.04157 | 4 | 166 |
| signal transduction involved in mitotic cell cycle checkpoint | 0.04157 | 3 | 68 |
| signal transduction involved in DNA damage checkpoint | 0.04157 | 3 | 68 |
| intracellular signal transduction involved in G1 DNA damage checkpoint | 0.04157 | 3 | 67 |
| signal transduction involved in cell cycle checkpoint | 0.041693 | 3 | 69 |
| positive regulation of ubiquitin-protein ligase activity involved in mitotic cell cycle | 0.041806 | 3 | 72 |
| antigen processing and presentation of exogenous peptide antigen via MHC class I, TAP-dependent | 0.041806 | 3 | 75 |
| mitotic G1 DNA damage checkpoint | 0.041806 | 3 | 74 |
| negative regulation of ubiquitin-protein ligase activity | 0.041806 | 3 | 71 |
| G1 DNA damage checkpoint | 0.041806 | 3 | 75 |
| negative regulation of ligase activity | 0.041806 | 3 | 71 |
| mitotic G1/S transition checkpoint | 0.041806 | 3 | 74 |
| antigen processing and presentation of exogenous peptide antigen via MHC class I | 0.044238 | 3 | 78 |
| regulation of ubiquitin-protein ligase activity involved in mitotic cell cycle | 0.044238 | 3 | 78 |
| 2-oxoglutarate metabolic process | 0.04747 | 2 | 13 |
| positive regulation of cell cycle arrest | 0.048104 | 3 | 83 |
| anaphase-promoting complex-dependent proteasomal ubiquitin-dependent protein catabolic process | 0.048104 | 3 | 83 |
| positive regulation of ubiquitin-protein ligase activity | 0.048104 | 3 | 84 |
| mitotic DNA damage checkpoint | 0.048104 | 3 | 84 |
| mitotic DNA integrity checkpoint | 0.050796 | 3 | 87 |
| positive regulation of ligase activity | 0.050796 | 3 | 87 |
| negative regulation of G1/S transition of mitotic cell cycle | 0.056677 | 3 | 91 |
| antigen processing and presentation of peptide antigen via MHC class I | 0.060955 | 3 | 94 |
| negative regulation of protein ubiquitination | 0.063409 | 3 | 96 |
| regulation of ubiquitin-protein ligase activity | 0.063935 | 3 | 97 |
| regulation of cell cycle arrest | 0.066996 | 3 | 100 |
| regulation of ligase activity | 0.066996 | 3 | 100 |
| DNA damage response, signal transduction by p53 class mediator | 0.084641 | 3 | 109 |
| regulation of G1/S transition of mitotic cell cycle | 0.087485 | 3 | 111 |
| negative regulation of cell cycle | 0.090561 | 4 | 290 |

**Table (5):**
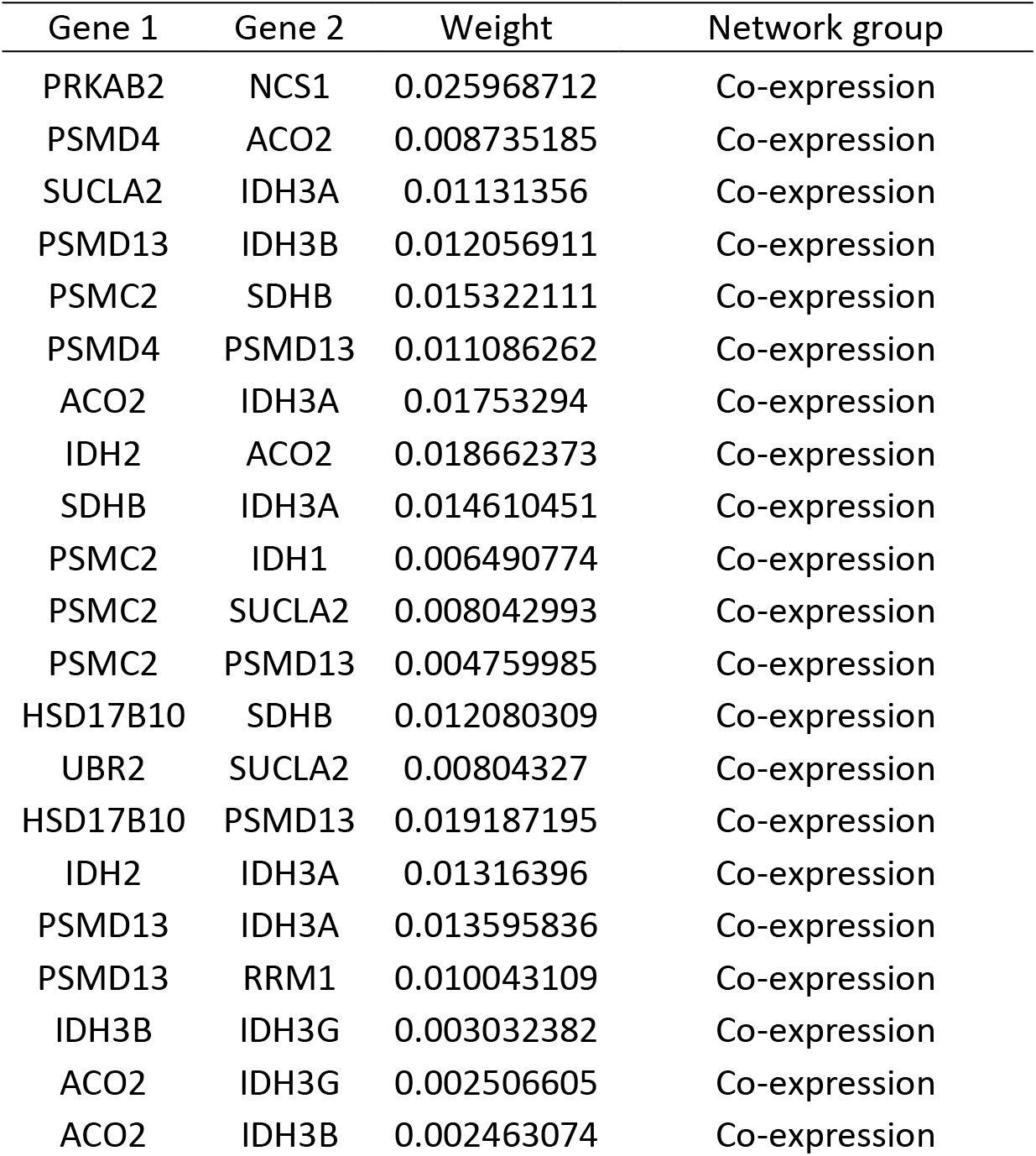
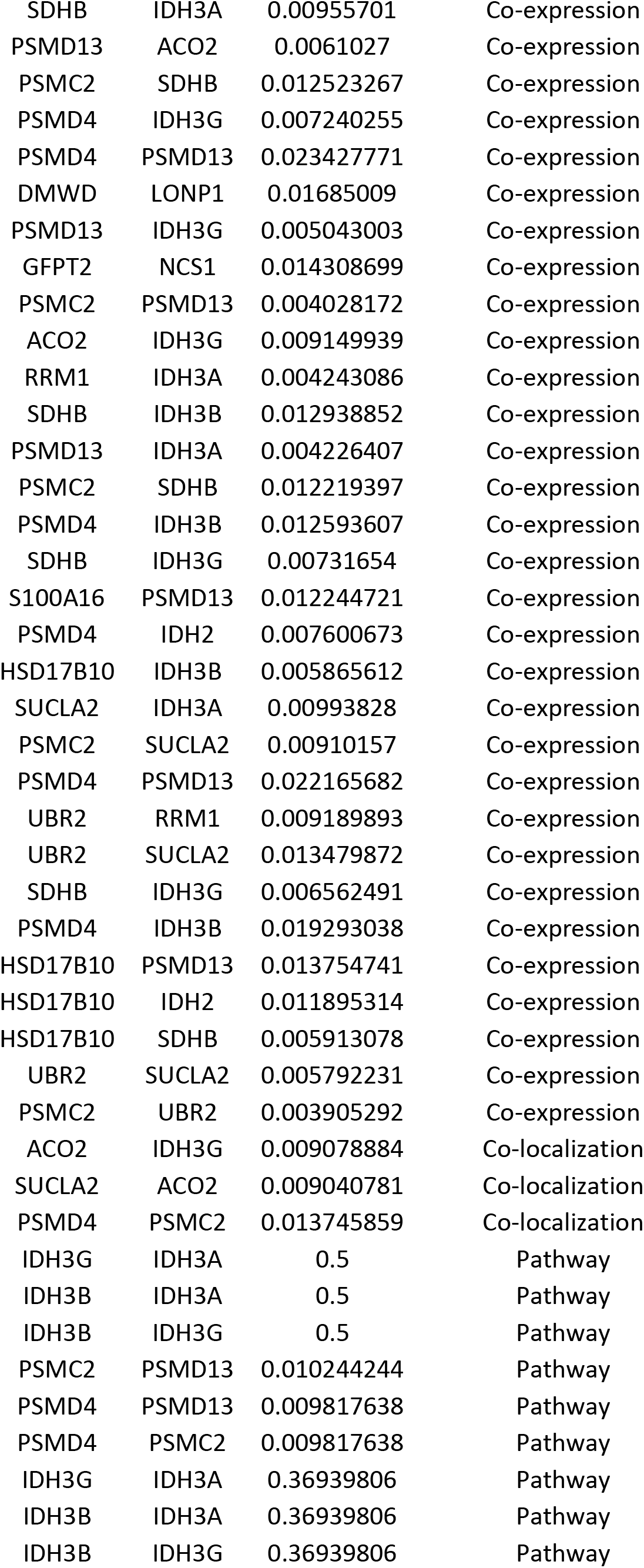
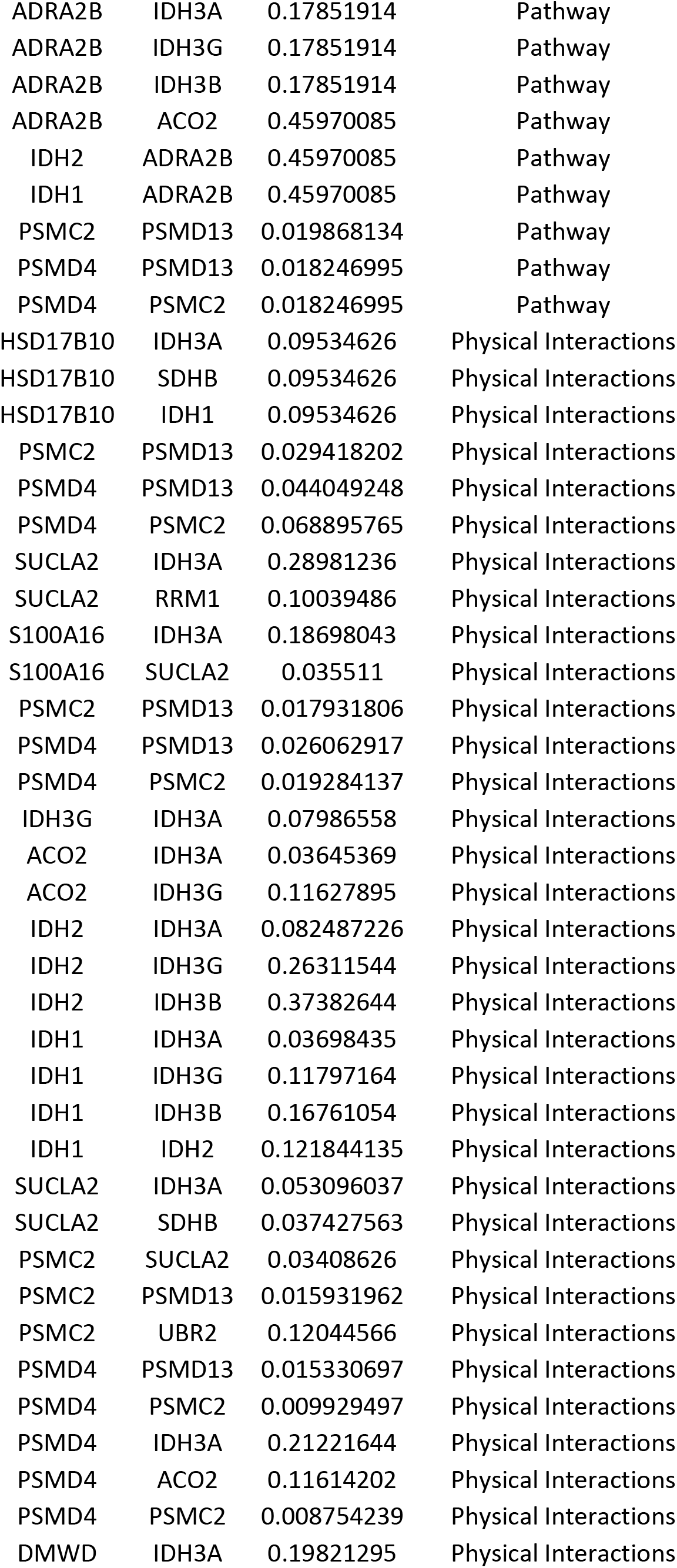
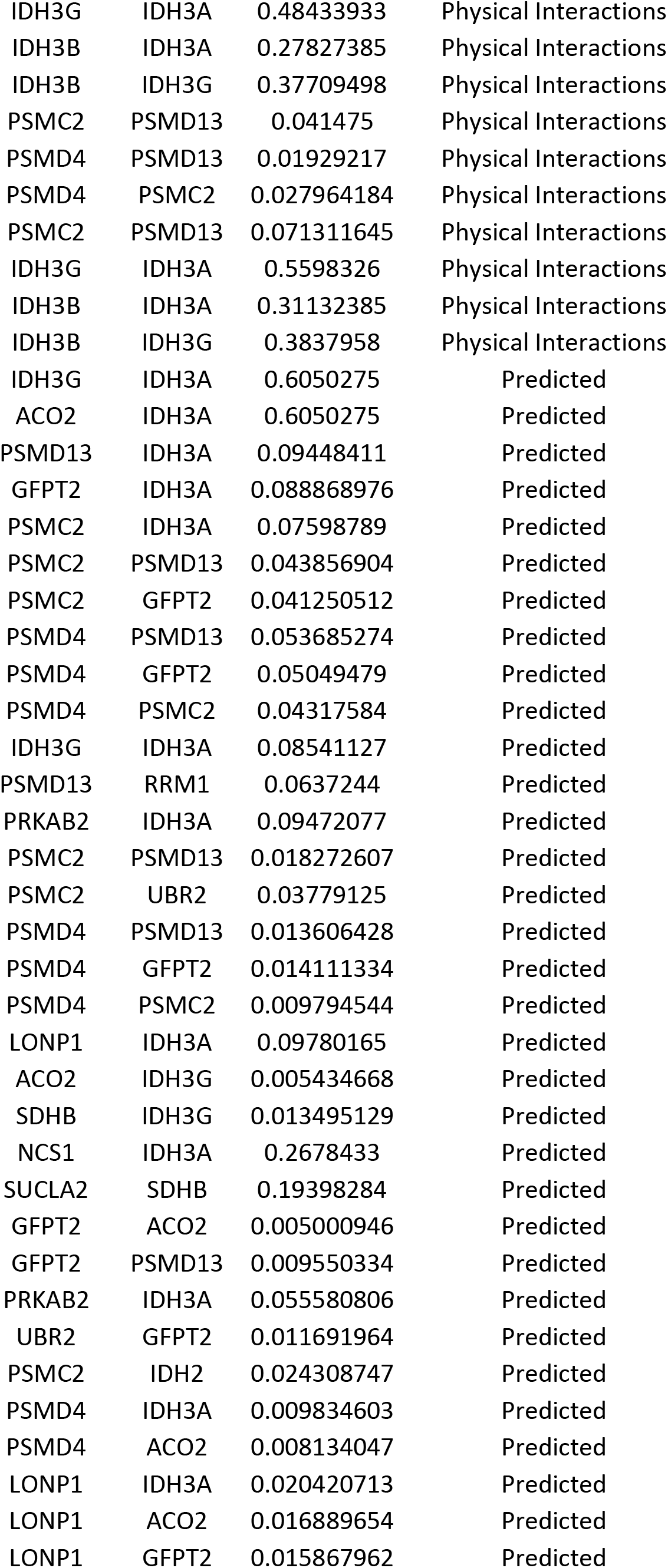
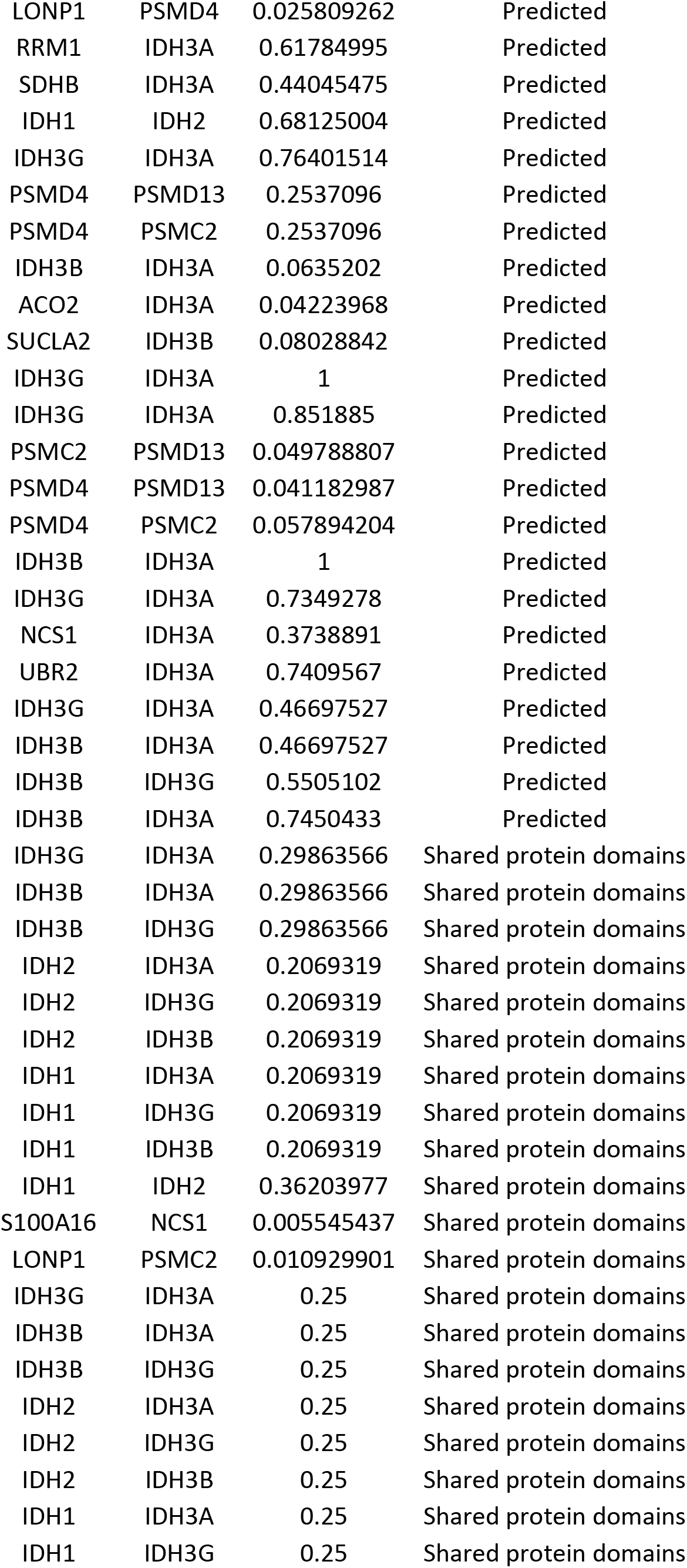
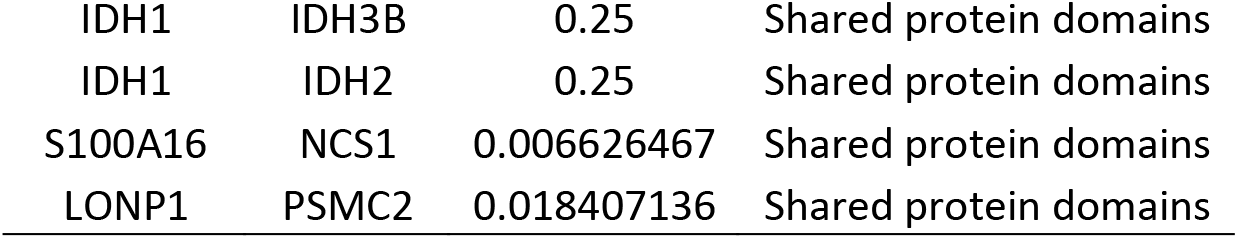
The gene co-expressed, shared domain and interaction with IDH3A gene network:

| Gene 1 | Gene 2 | Weight | Network group |
| --- | --- | --- | --- |
| PRKAB2 | NCS1 | 0.025968712 | Co-expression |
| PSMD4 | ACO2 | 0.008735185 | Co-expression |
| SUCLA2 | IDH3A | 0.01131356 | Co-expression |
| PSMD13 | IDH3B | 0.012056911 | Co-expression |
| PSMC2 | SDHB | 0.015322111 | Co-expression |
| PSMD4 | PSMD13 | 0.011086262 | Co-expression |
| ACO2 | IDH3A | 0.01753294 | Co-expression |
| IDH2 | ACO2 | 0.018662373 | Co-expression |
| SDHB | IDH3A | 0.014610451 | Co-expression |
| PSMC2 | IDH1 | 0.006490774 | Co-expression |
| PSMC2 | SUCLA2 | 0.008042993 | Co-expression |
| PSMC2 | PSMD13 | 0.004759985 | Co-expression |
| HSD17B10 | SDHB | 0.012080309 | Co-expression |
| UBR2 | SUCLA2 | 0.00804327 | Co-expression |
| HSD17B10 | PSMD13 | 0.019187195 | Co-expression |
| IDH2 | IDH3A | 0.01316396 | Co-expression |
| PSMD13 | IDH3A | 0.013595836 | Co-expression |
| PSMD13 | RRM1 | 0.010043109 | Co-expression |
| IDH3B | IDH3G | 0.003032382 | Co-expression |
| ACO2 | IDH3G | 0.002506605 | Co-expression |
| ACO2 | IDH3B | 0.002463074 | Co-expression |
| SDHB | IDH3A | 0.00955701 | Co-expression |
| PSMD13 | ACO2 | 0.0061027 | Co-expression |
| PSMC2 | SDHB | 0.012523267 | Co-expression |
| PSMD4 | IDH3G | 0.007240255 | Co-expression |
| PSMD4 | PSMD13 | 0.023427771 | Co-expression |
| DMWD | LONP1 | 0.01685009 | Co-expression |
| PSMD13 | IDH3G | 0.005043003 | Co-expression |
| GFPT2 | NCS1 | 0.014308699 | Co-expression |
| PSMC2 | PSMD13 | 0.004028172 | Co-expression |
| ACO2 | IDH3G | 0.009149939 | Co-expression |
| RRM1 | IDH3A | 0.004243086 | Co-expression |
| SDHB | IDH3B | 0.012938852 | Co-expression |
| PSMD13 | IDH3A | 0.004226407 | Co-expression |
| PSMC2 | SDHB | 0.012219397 | Co-expression |
| PSMD4 | IDH3B | 0.012593607 | Co-expression |
| SDHB | IDH3G | 0.00731654 | Co-expression |
| S100A16 | PSMD13 | 0.012244721 | Co-expression |
| PSMD4 | IDH2 | 0.007600673 | Co-expression |
| HSD17B10 | IDH3B | 0.005865612 | Co-expression |
| SUCLA2 | IDH3A | 0.00993828 | Co-expression |
| PSMC2 | SUCLA2 | 0.00910157 | Co-expression |
| PSMD4 | PSMD13 | 0.022165682 | Co-expression |
| UBR2 | RRM1 | 0.009189893 | Co-expression |
| UBR2 | SUCLA2 | 0.013479872 | Co-expression |
| SDHB | IDH3G | 0.006562491 | Co-expression |
| PSMD4 | IDH3B | 0.019293038 | Co-expression |
| HSD17B10 | PSMD13 | 0.013754741 | Co-expression |
| HSD17B10 | IDH2 | 0.011895314 | Co-expression |
| HSD17B10 | SDHB | 0.005913078 | Co-expression |
| UBR2 | SUCLA2 | 0.005792231 | Co-expression |
| PSMC2 | UBR2 | 0.003905292 | Co-expression |
| ACO2 | IDH3G | 0.009078884 | Co-localization |
| SUCLA2 | ACO2 | 0.009040781 | Co-localization |
| PSMD4 | PSMC2 | 0.013745859 | Co-localization |
| IDH3G | IDH3A | 0.5 | Pathway |
| IDH3B | IDH3A | 0.5 | Pathway |
| IDH3B | IDH3G | 0.5 | Pathway |
| PSMC2 | PSMD13 | 0.010244244 | Pathway |
| PSMD4 | PSMD13 | 0.009817638 | Pathway |
| PSMD4 | PSMC2 | 0.009817638 | Pathway |
| IDH3G | IDH3A | 0.36939806 | Pathway |
| IDH3B | IDH3A | 0.36939806 | Pathway |
| IDH3B | IDH3G | 0.36939806 | Pathway |
| ADRA2B | IDH3A | 0.17851914 | Pathway |
| ADRA2B | IDH3G | 0.17851914 | Pathway |
| ADRA2B | IDH3B | 0.17851914 | Pathway |
| ADRA2B | ACO2 | 0.45970085 | Pathway |
| IDH2 | ADRA2B | 0.45970085 | Pathway |
| IDH1 | ADRA2B | 0.45970085 | Pathway |
| PSMC2 | PSMD13 | 0.019868134 | Pathway |
| PSMD4 | PSMD13 | 0.018246995 | Pathway |
| PSMD4 | PSMC2 | 0.018246995 | Pathway |
| HSD17B10 | IDH3A | 0.09534626 | Physical Interactions |
| HSD17B10 | SDHB | 0.09534626 | Physical Interactions |
| HSD17B10 | IDH1 | 0.09534626 | Physical Interactions |
| PSMC2 | PSMD13 | 0.029418202 | Physical Interactions |
| PSMD4 | PSMD13 | 0.044049248 | Physical Interactions |
| PSMD4 | PSMC2 | 0.068895765 | Physical Interactions |
| SUCLA2 | IDH3A | 0.28981236 | Physical Interactions |
| SUCLA2 | RRM1 | 0.10039486 | Physical Interactions |
| S100A16 | IDH3A | 0.18698043 | Physical Interactions |
| S100A16 | SUCLA2 | 0.035511 | Physical Interactions |
| PSMC2 | PSMD13 | 0.017931806 | Physical Interactions |
| PSMD4 | PSMD13 | 0.026062917 | Physical Interactions |
| PSMD4 | PSMC2 | 0.019284137 | Physical Interactions |
| IDH3G | IDH3A | 0.07986558 | Physical Interactions |
| ACO2 | IDH3A | 0.03645369 | Physical Interactions |
| ACO2 | IDH3G | 0.11627895 | Physical Interactions |
| IDH2 | IDH3A | 0.082487226 | Physical Interactions |
| IDH2 | IDH3G | 0.26311544 | Physical Interactions |
| IDH2 | IDH3B | 0.37382644 | Physical Interactions |
| IDH1 | IDH3A | 0.03698435 | Physical Interactions |
| IDH1 | IDH3G | 0.11797164 | Physical Interactions |
| IDH1 | IDH3B | 0.16761054 | Physical Interactions |
| IDH1 | IDH2 | 0.121844135 | Physical Interactions |
| SUCLA2 | IDH3A | 0.053096037 | Physical Interactions |
| SUCLA2 | SDHB | 0.037427563 | Physical Interactions |
| PSMC2 | SUCLA2 | 0.03408626 | Physical Interactions |
| PSMC2 | PSMD13 | 0.015931962 | Physical Interactions |
| PSMC2 | UBR2 | 0.12044566 | Physical Interactions |
| PSMD4 | PSMD13 | 0.015330697 | Physical Interactions |
| PSMD4 | PSMC2 | 0.009929497 | Physical Interactions |
| PSMD4 | IDH3A | 0.21221644 | Physical Interactions |
| PSMD4 | ACO2 | 0.11614202 | Physical Interactions |
| PSMD4 | PSMC2 | 0.008754239 | Physical Interactions |
| DMWD | IDH3A | 0.19821295 | Physical Interactions |
| IDH3G | IDH3A | 0.48433933 | Physical Interactions |
| IDH3B | IDH3A | 0.27827385 | Physical Interactions |
| IDH3B | IDH3G | 0.37709498 | Physical Interactions |
| PSMC2 | PSMD13 | 0.041475 | Physical Interactions |
| PSMD4 | PSMD13 | 0.01929217 | Physical Interactions |
| PSMD4 | PSMC2 | 0.027964184 | Physical Interactions |
| PSMC2 | PSMD13 | 0.071311645 | Physical Interactions |
| IDH3G | IDH3A | 0.5598326 | Physical Interactions |
| IDH3B | IDH3A | 0.31132385 | Physical Interactions |
| IDH3B | IDH3G | 0.3837958 | Physical Interactions |
| IDH3G | IDH3A | 0.6050275 | Predicted |
| ACO2 | IDH3A | 0.6050275 | Predicted |
| PSMD13 | IDH3A | 0.09448411 | Predicted |
| GFPT2 | IDH3A | 0.088868976 | Predicted |
| PSMC2 | IDH3A | 0.07598789 | Predicted |
| PSMC2 | PSMD13 | 0.043856904 | Predicted |
| PSMC2 | GFPT2 | 0.041250512 | Predicted |
| PSMD4 | PSMD13 | 0.053685274 | Predicted |
| PSMD4 | GFPT2 | 0.05049479 | Predicted |
| PSMD4 | PSMC2 | 0.04317584 | Predicted |
| IDH3G | IDH3A | 0.08541127 | Predicted |
| PSMD13 | RRM1 | 0.0637244 | Predicted |
| PRKAB2 | IDH3A | 0.09472077 | Predicted |
| PSMC2 | PSMD13 | 0.018272607 | Predicted |
| PSMC2 | UBR2 | 0.03779125 | Predicted |
| PSMD4 | PSMD13 | 0.013606428 | Predicted |
| PSMD4 | GFPT2 | 0.014111334 | Predicted |
| PSMD4 | PSMC2 | 0.009794544 | Predicted |
| LONP1 | IDH3A | 0.09780165 | Predicted |
| ACO2 | IDH3G | 0.005434668 | Predicted |
| SDHB | IDH3G | 0.013495129 | Predicted |
| NCS1 | IDH3A | 0.2678433 | Predicted |
| SUCLA2 | SDHB | 0.19398284 | Predicted |
| GFPT2 | ACO2 | 0.005000946 | Predicted |
| GFPT2 | PSMD13 | 0.009550334 | Predicted |
| PRKAB2 | IDH3A | 0.055580806 | Predicted |
| UBR2 | GFPT2 | 0.011691964 | Predicted |
| PSMC2 | IDH2 | 0.024308747 | Predicted |
| PSMD4 | IDH3A | 0.009834603 | Predicted |
| PSMD4 | ACO2 | 0.008134047 | Predicted |
| LONP1 | IDH3A | 0.020420713 | Predicted |
| LONP1 | ACO2 | 0.016889654 | Predicted |
| LONP1 | GFPT2 | 0.015867962 | Predicted |
| LONP1 | PSMD4 | 0.025809262 | Predicted |
| RRM1 | IDH3A | 0.61784995 | Predicted |
| SDHB | IDH3A | 0.44045475 | Predicted |
| IDH1 | IDH2 | 0.68125004 | Predicted |
| IDH3G | IDH3A | 0.76401514 | Predicted |
| PSMD4 | PSMD13 | 0.2537096 | Predicted |
| PSMD4 | PSMC2 | 0.2537096 | Predicted |
| IDH3B | IDH3A | 0.0635202 | Predicted |
| ACO2 | IDH3A | 0.04223968 | Predicted |
| SUCLA2 | IDH3B | 0.08028842 | Predicted |
| IDH3G | IDH3A | 1 | Predicted |
| IDH3G | IDH3A | 0.851885 | Predicted |
| PSMC2 | PSMD13 | 0.049788807 | Predicted |
| PSMD4 | PSMD13 | 0.041182987 | Predicted |
| PSMD4 | PSMC2 | 0.057894204 | Predicted |
| IDH3B | IDH3A | 1 | Predicted |
| IDH3G | IDH3A | 0.7349278 | Predicted |
| NCS1 | IDH3A | 0.3738891 | Predicted |
| UBR2 | IDH3A | 0.7409567 | Predicted |
| IDH3G | IDH3A | 0.46697527 | Predicted |
| IDH3B | IDH3A | 0.46697527 | Predicted |
| IDH3B | IDH3G | 0.5505102 | Predicted |
| IDH3B | IDH3A | 0.7450433 | Predicted |
| IDH3G | IDH3A | 0.29863566 | Shared protein domains |
| IDH3B | IDH3A | 0.29863566 | Shared protein domains |
| IDH3B | IDH3G | 0.29863566 | Shared protein domains |
| IDH2 | IDH3A | 0.2069319 | Shared protein domains |
| IDH2 | IDH3G | 0.2069319 | Shared protein domains |
| IDH2 | IDH3B | 0.2069319 | Shared protein domains |
| IDH1 | IDH3A | 0.2069319 | Shared protein domains |
| IDH1 | IDH3G | 0.2069319 | Shared protein domains |
| IDH1 | IDH3B | 0.2069319 | Shared protein domains |
| IDH1 | IDH2 | 0.36203977 | Shared protein domains |
| S100A16 | NCS1 | 0.005545437 | Shared protein domains |
| LONP1 | PSMC2 | 0.010929901 | Shared protein domains |
| IDH3G | IDH3A | 0.25 | Shared protein domains |
| IDH3B | IDH3A | 0.25 | Shared protein domains |
| IDH3B | IDH3G | 0.25 | Shared protein domains |
| IDH2 | IDH3A | 0.25 | Shared protein domains |
| IDH2 | IDH3G | 0.25 | Shared protein domains |
| IDH2 | IDH3B | 0.25 | Shared protein domains |
| IDH1 | IDH3A | 0.25 | Shared protein domains |
| IDH1 | IDH3G | 0.25 | Shared protein domains |
| IDH1 | IDH3B | 0.25 | Shared protein domains |
| IDH1 | IDH2 | 0.25 | Shared protein domains |
| S100A16 | NCS1 | 0.006626467 | Shared protein domains |
| LONP1 | PSMC2 | 0.018407136 | Shared protein domains |

**Figure 1:**
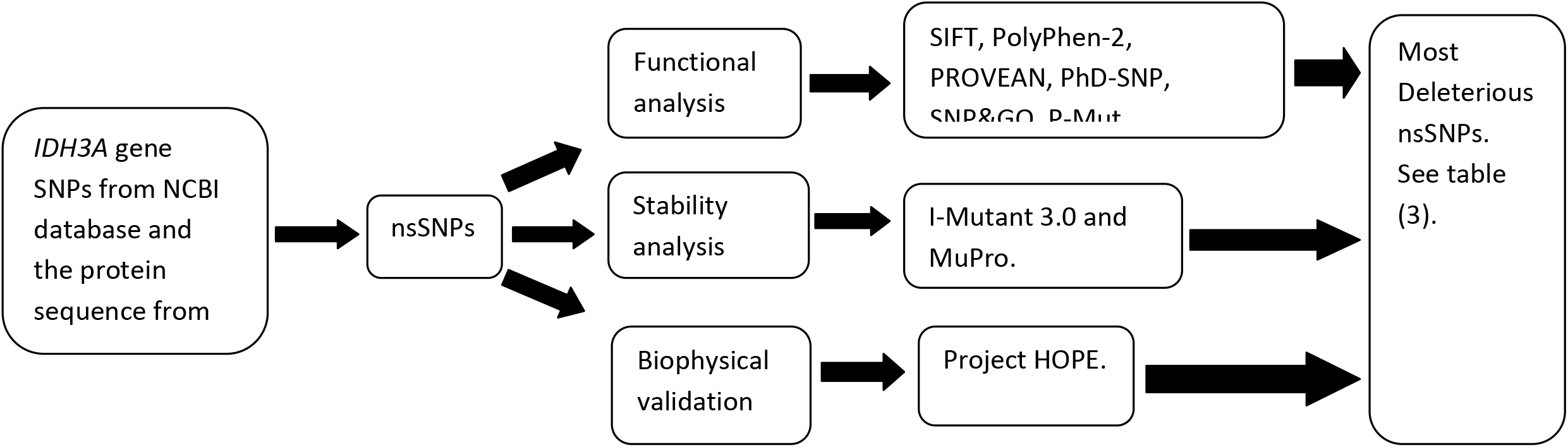
Diagrammatic representation of the *IDH3A* gene in silico work flow.

**Figure 2:**
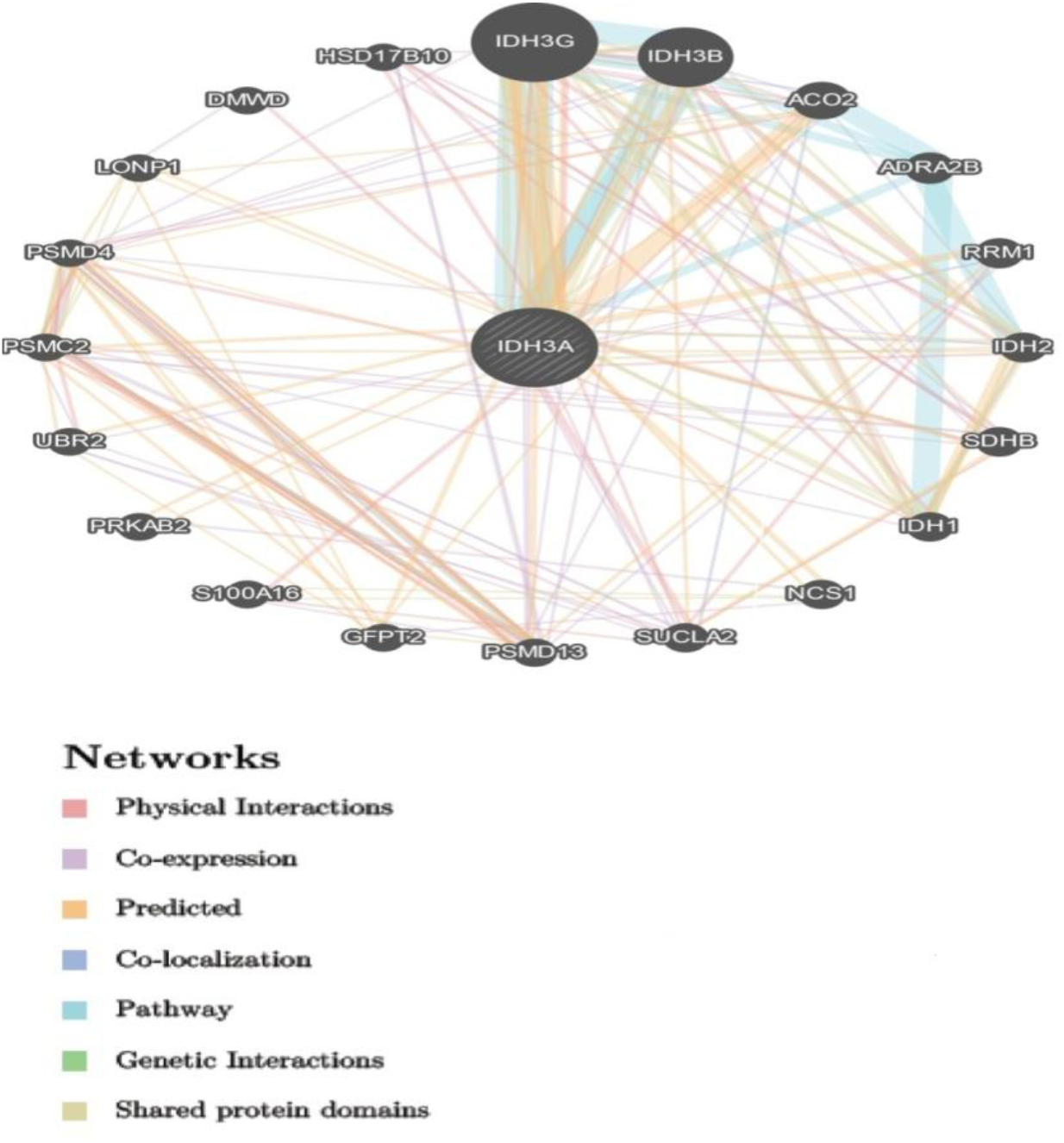
Interaction of *IDH3A* gene with its related genes.

**Figure 3:**
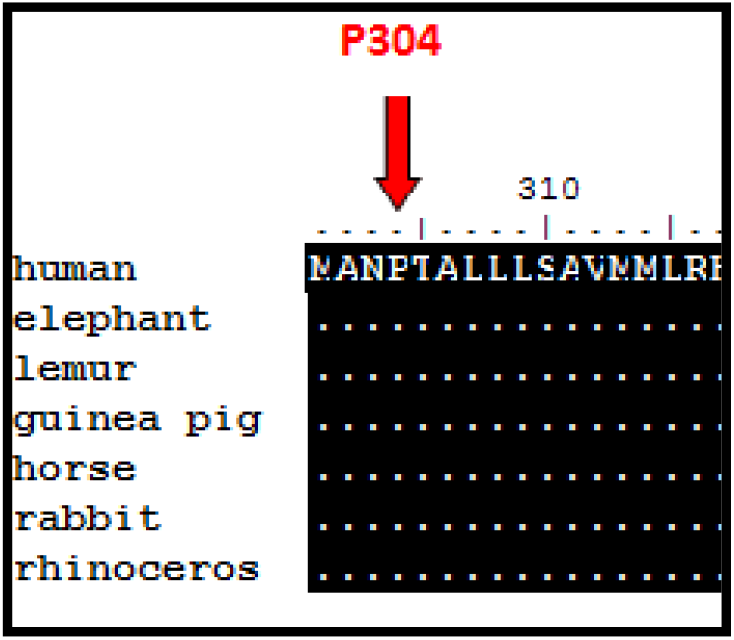
Alignment of 7 amino acid sequences of *IDH3A* gene shows that the residue predicted to be mutated in our study (pointed at by a red arrow) is actually found in a conserved region across species. Sequence alignment was done using BioEdit software (v7.2.5).

**Figure 4:**
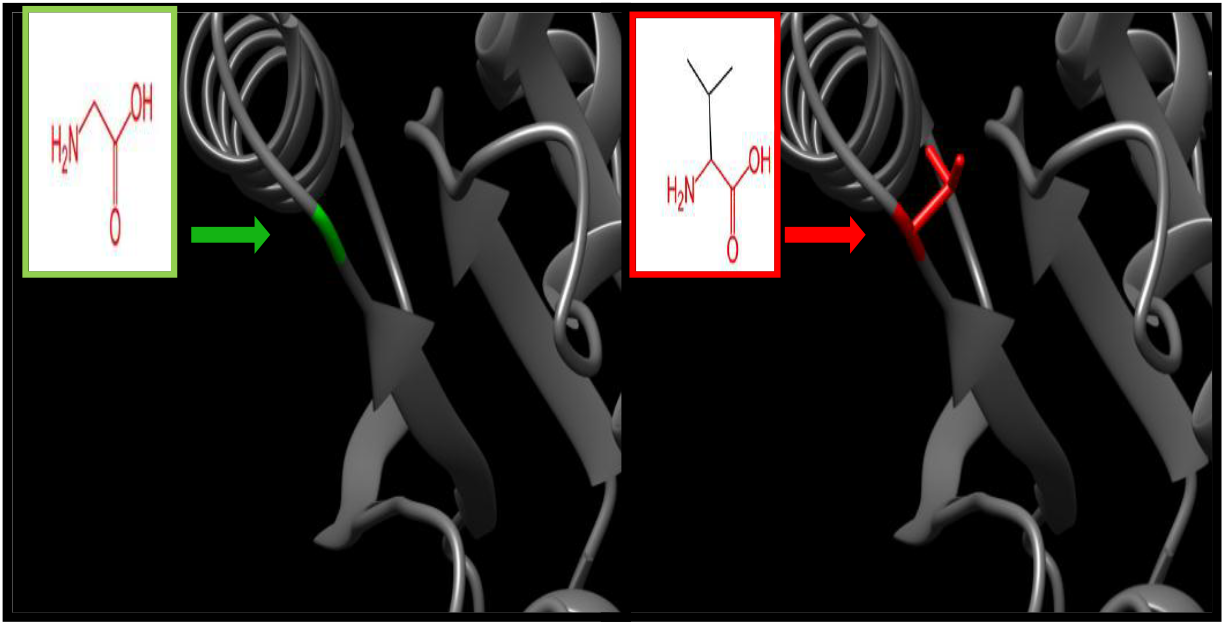
(rs1465252689) (G39V) the amino acid Glycine changes to Valine at position 39.

**Figure 5:**
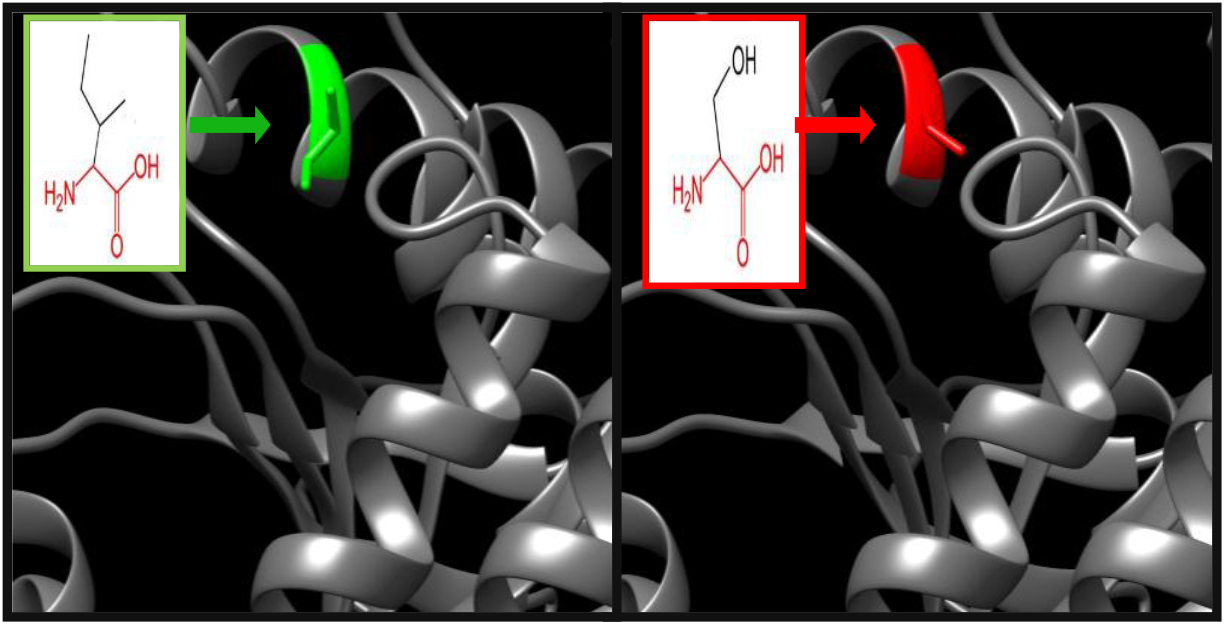
(rs1271001430) (I46S) the amino acid Isoleucine changes to Serine at position 46.

**Figure 6:**
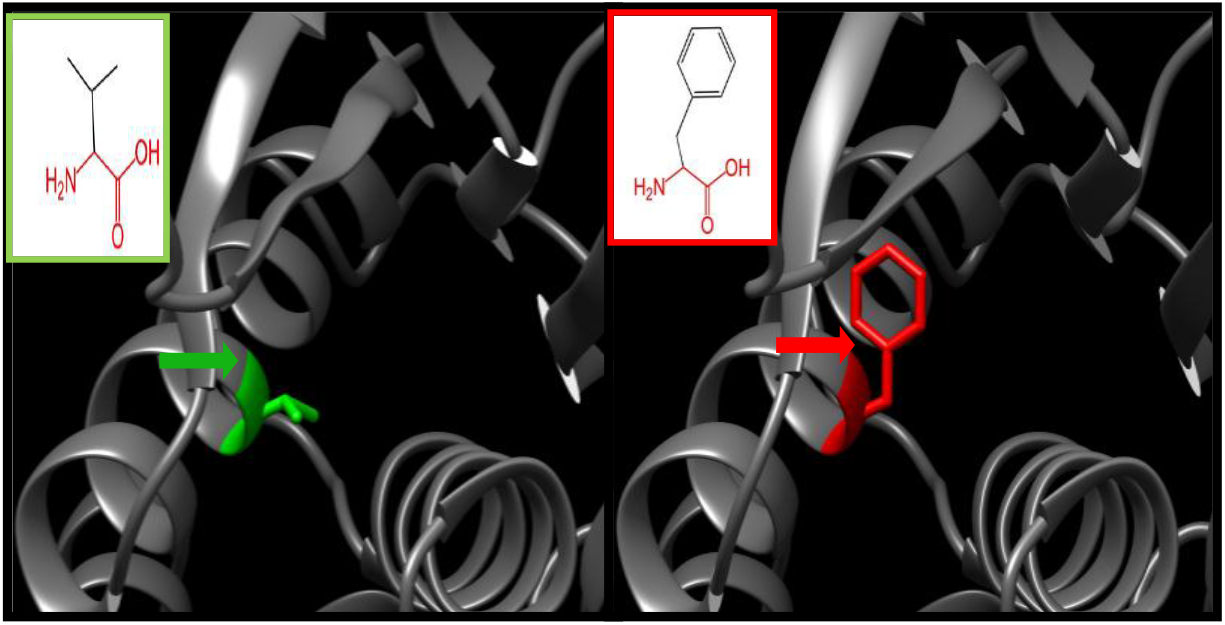
(V50F) the amino acid Valine changes to Phenylalanine at position 50.

**Figure 7:**
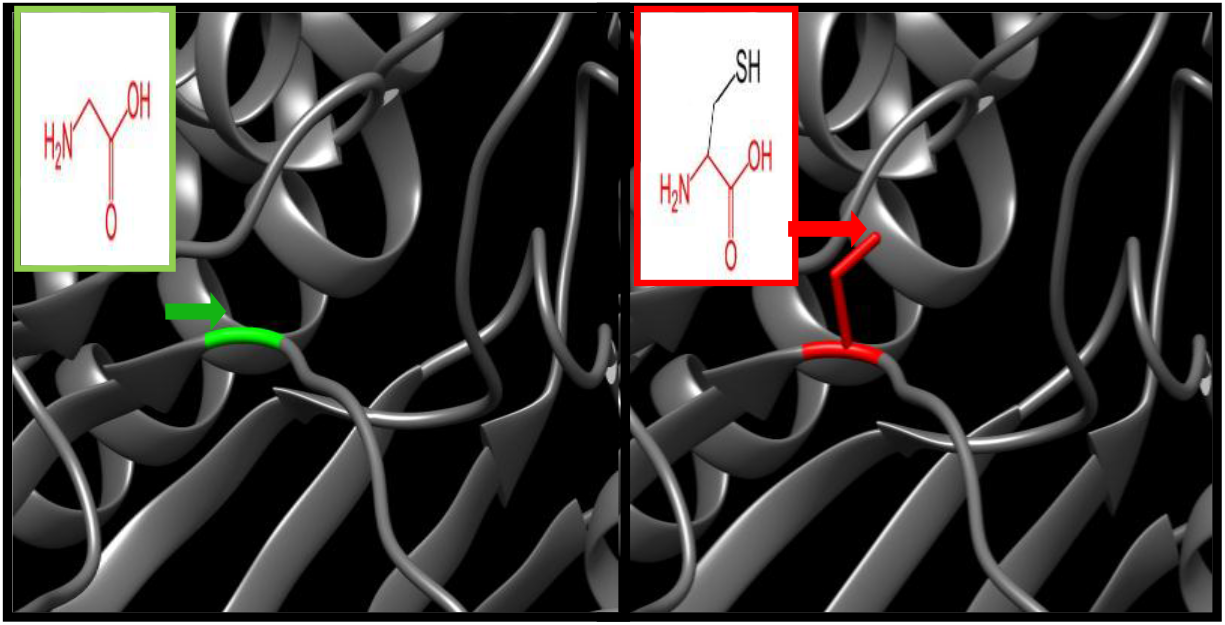
(rs1205294920) (G97C) the amino acid Glycine changes to Cysteine at position 97.

**Figure 8:**
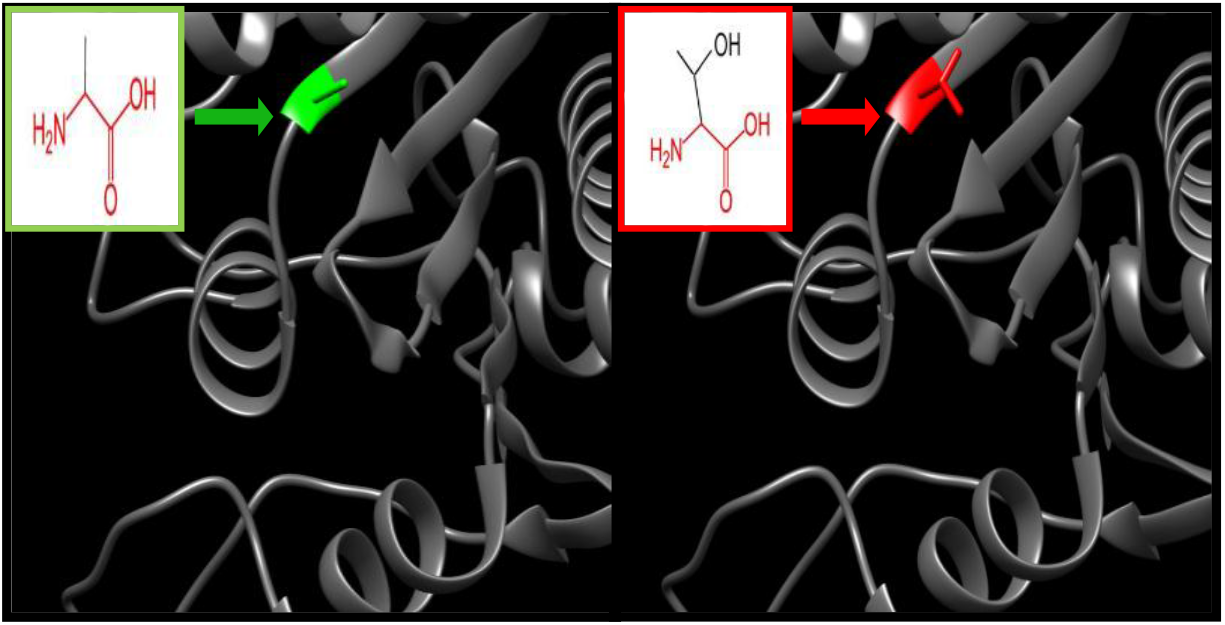
(rs756333430) (A122T) the amino acid Alanine changes to Threonine at position 122.

**Figure 9:**
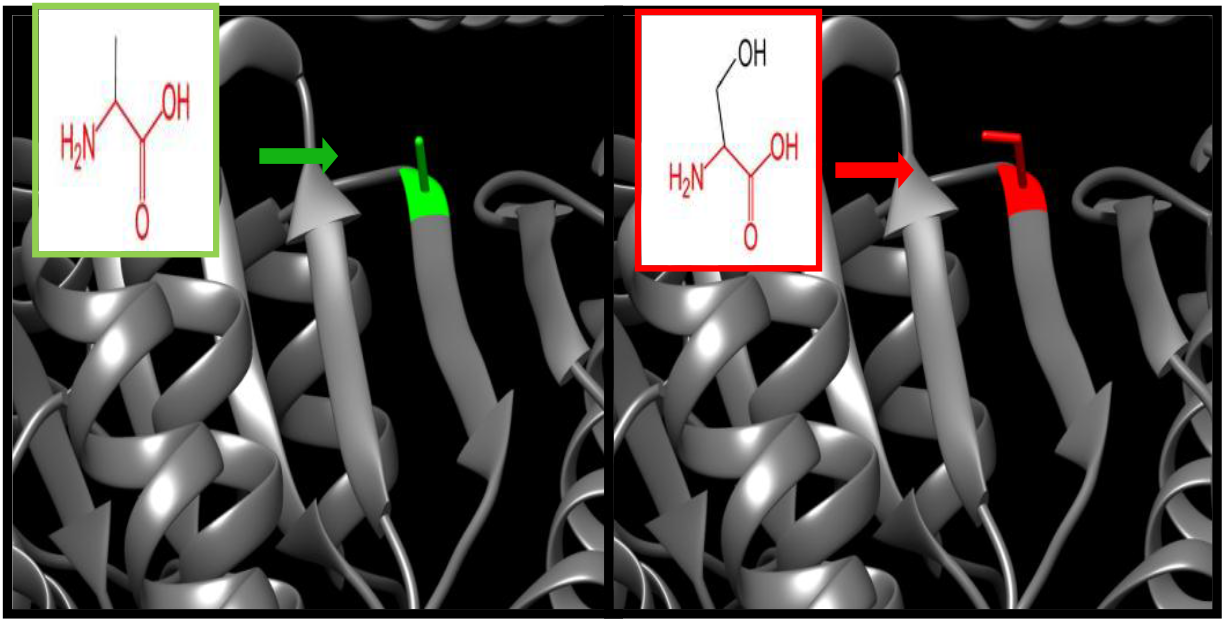
(A122S) the amino acid Alanine changes to Serine at position 122.

**Figure 10:**
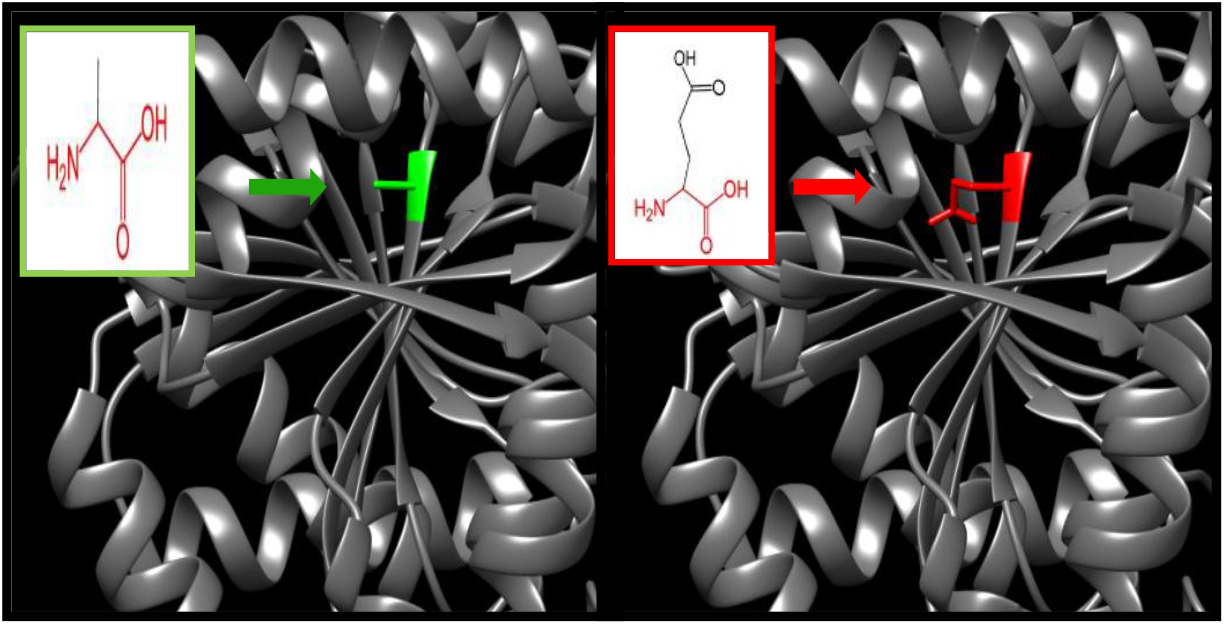
(rs766657715) (A122E) the amino acid Alanine changes to Glutamic Acid at position 122.

**Figure 11:**
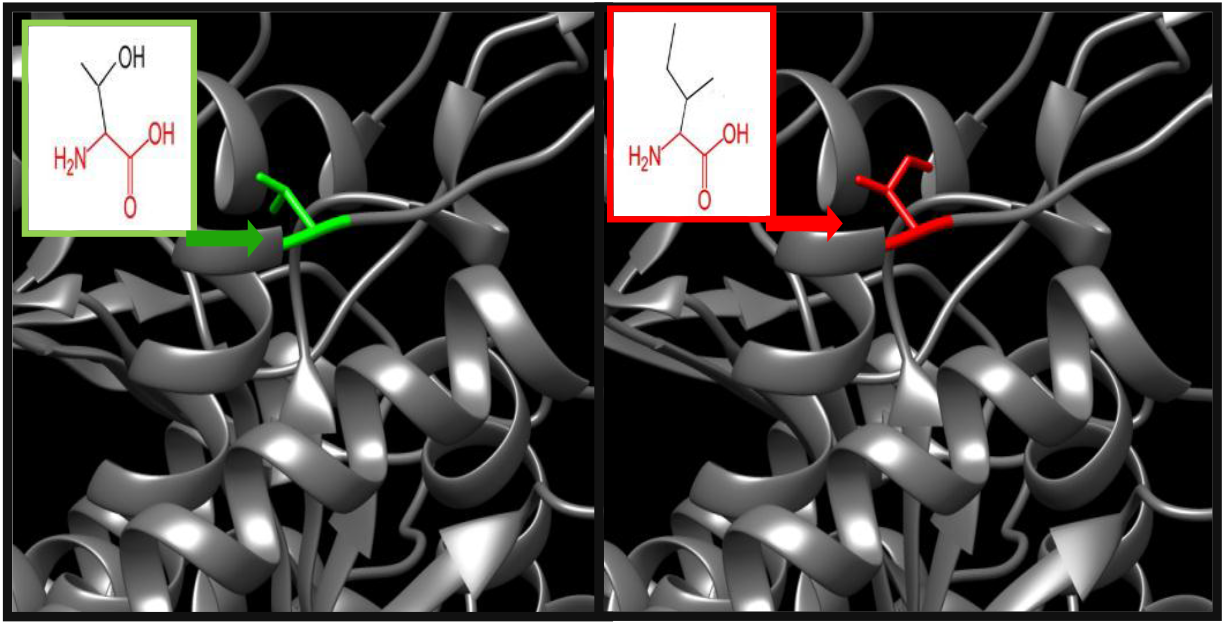
(rs1257538190) (T172I) the amino acid Threonine changes to Isoleucine at position 172.

**Figure 12:**
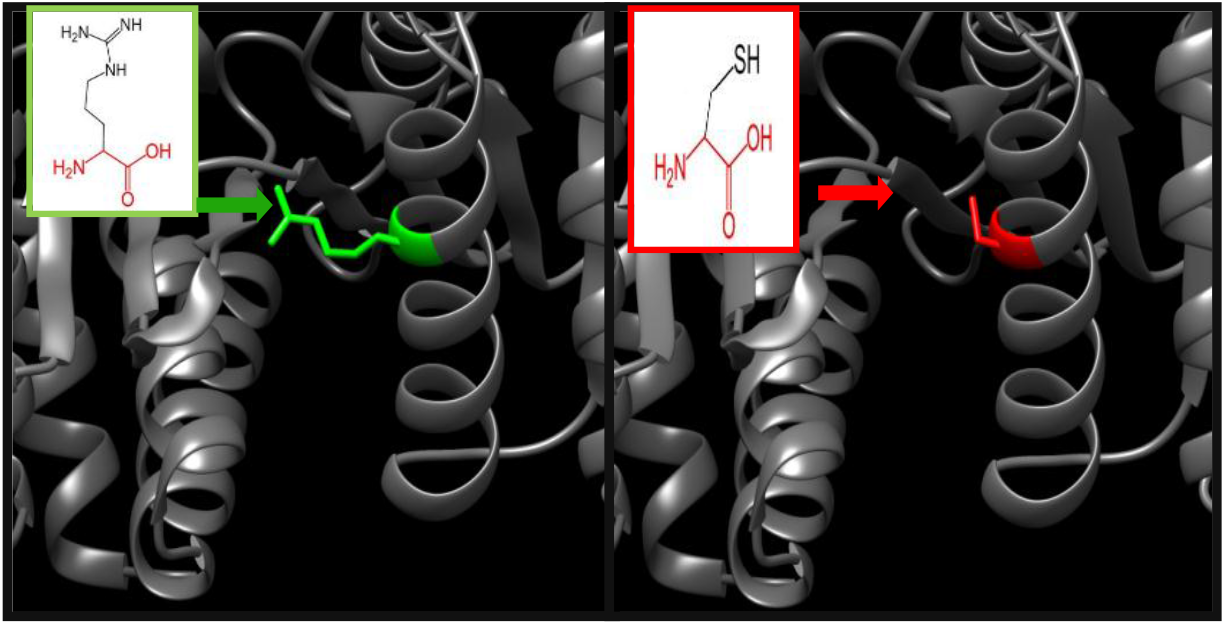
(rs1404589314) (R178C) the amino acid Arginine changes to Cysteine at position 178.

**Figure 13:**
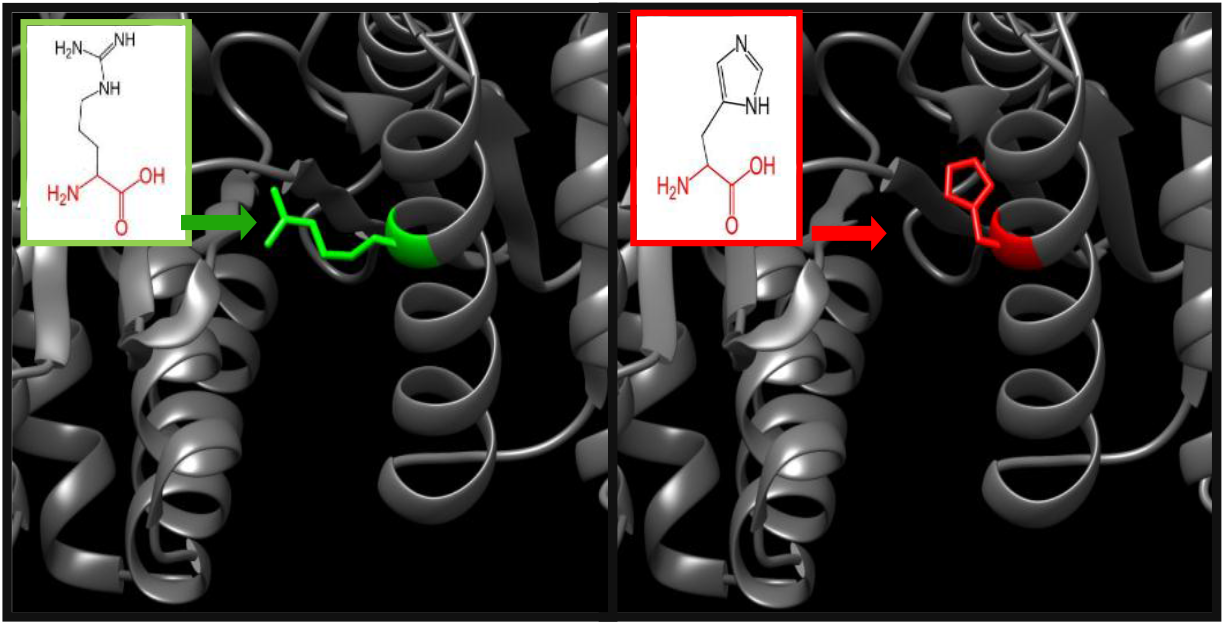
(rs1329563873) (R178H) the amino acid Arginine changes to Histidine at position 178.

**Figure 14:**
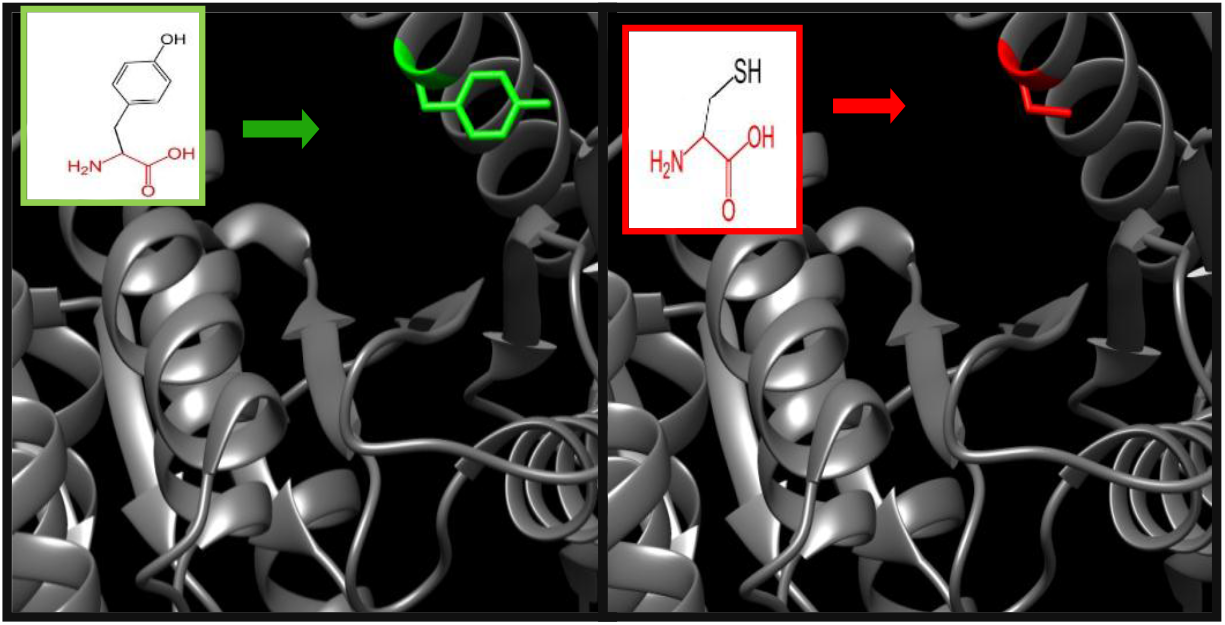
(rs1303775777) (Y186C) the amino acid Tyrosine changes to Cysteine at position 186.

**Figure 15:**
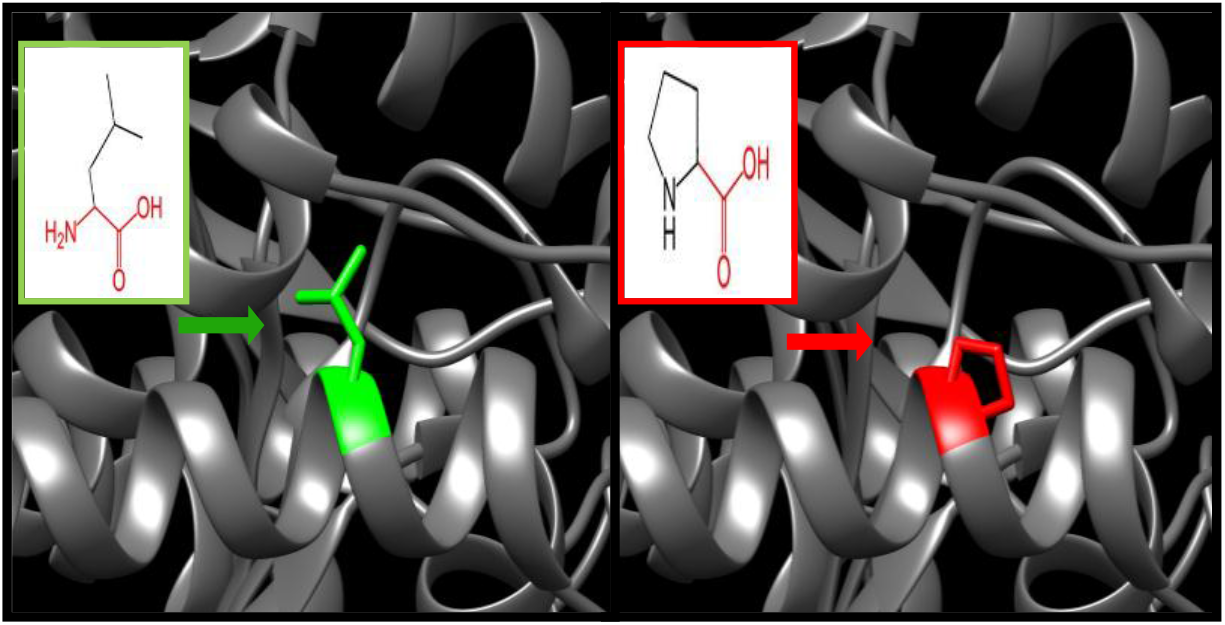
(rs200797531) (V198G) the amino acid Valine changes to Glycine at position 198.

**Figure 16:**
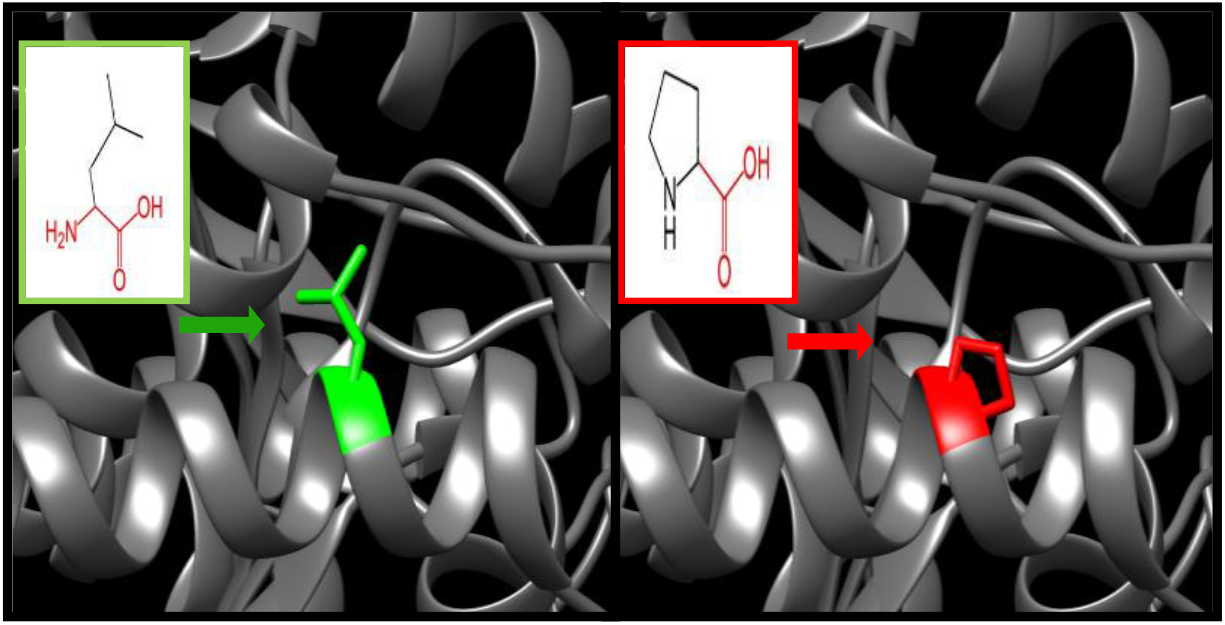
(rs745664361) (L210P) the amino acid Leucine changes to Proline at position 210.

**Figure 17:**
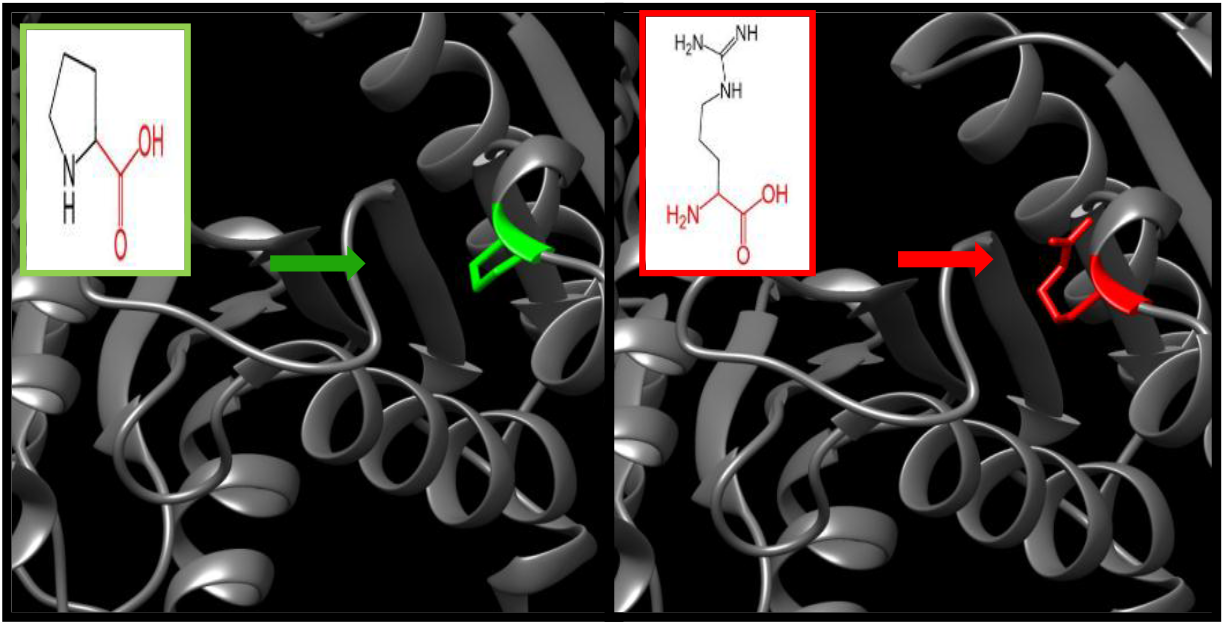
(rs978588480) (P243R) the amino acid Proline changes to Arginine at position 243.

**Figure 18:**
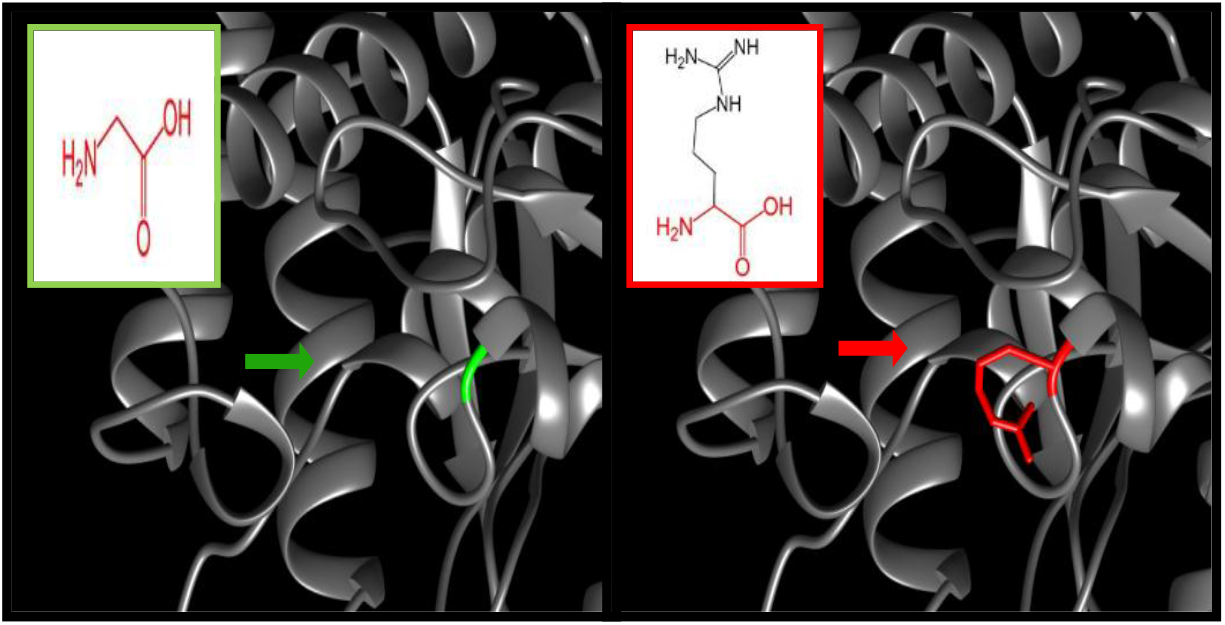
(rs1429951757) (G268R) the amino acid Glycine changes to Arginine at position 268.

**Figure 19:**
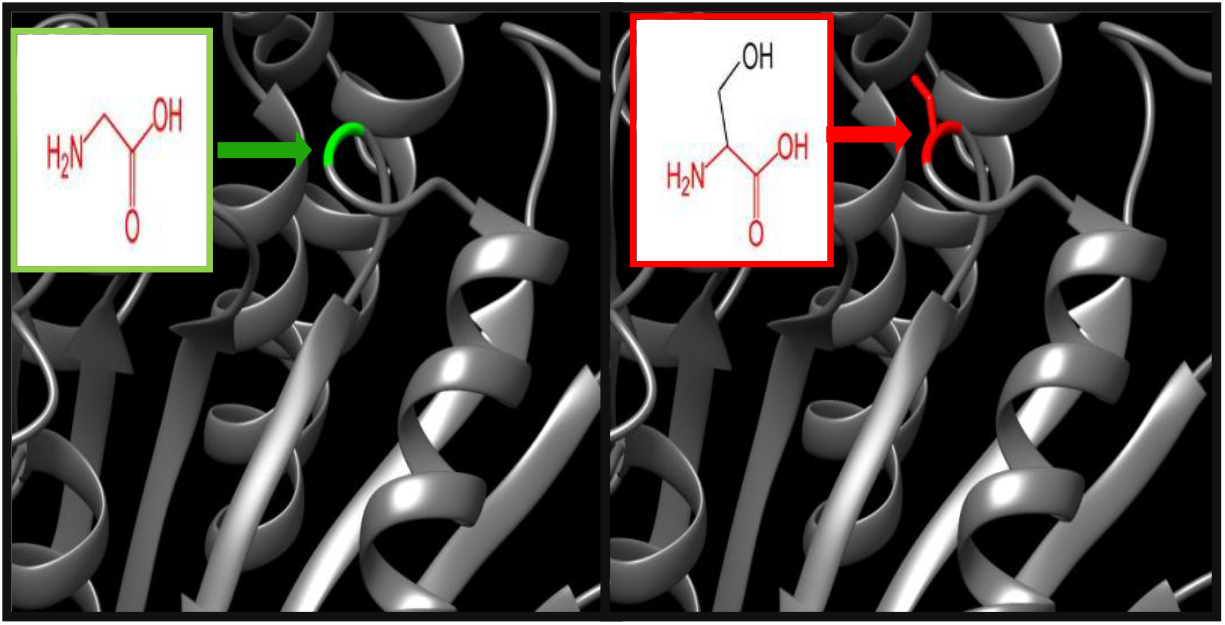
(rs775992590) (G271S) the amino acid Glycine changes to Serine at position 271.

**Figure 20:**
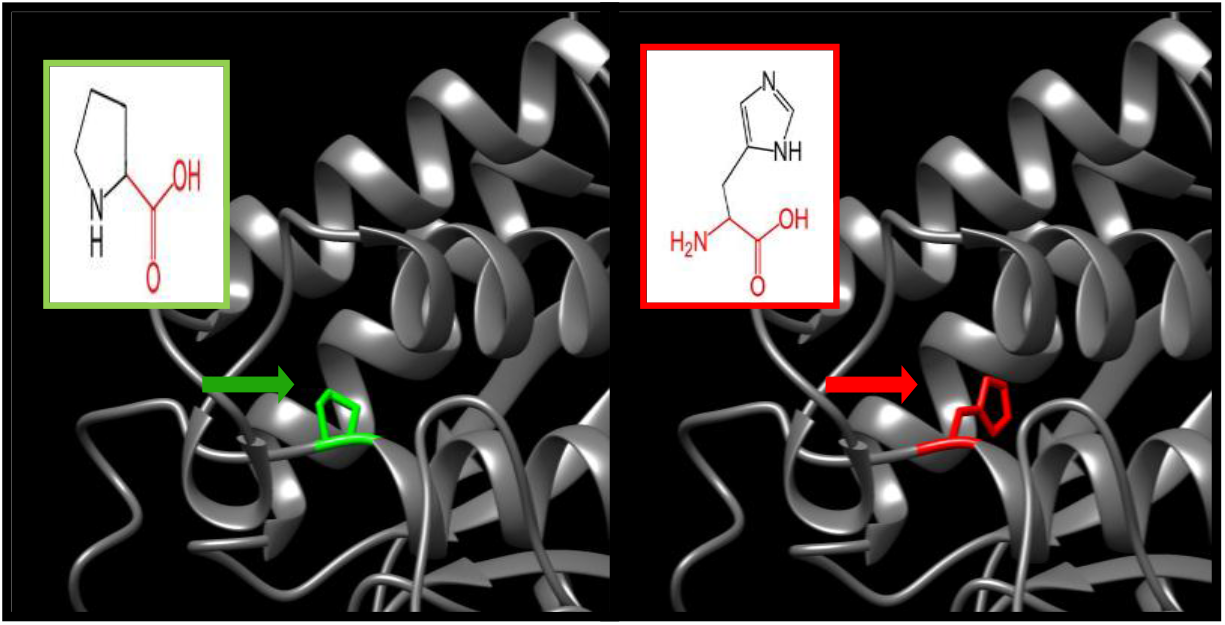
(rs756712426) (P304H) the amino acid Proline changes to Histidine at position 304.

**Figure 21:**
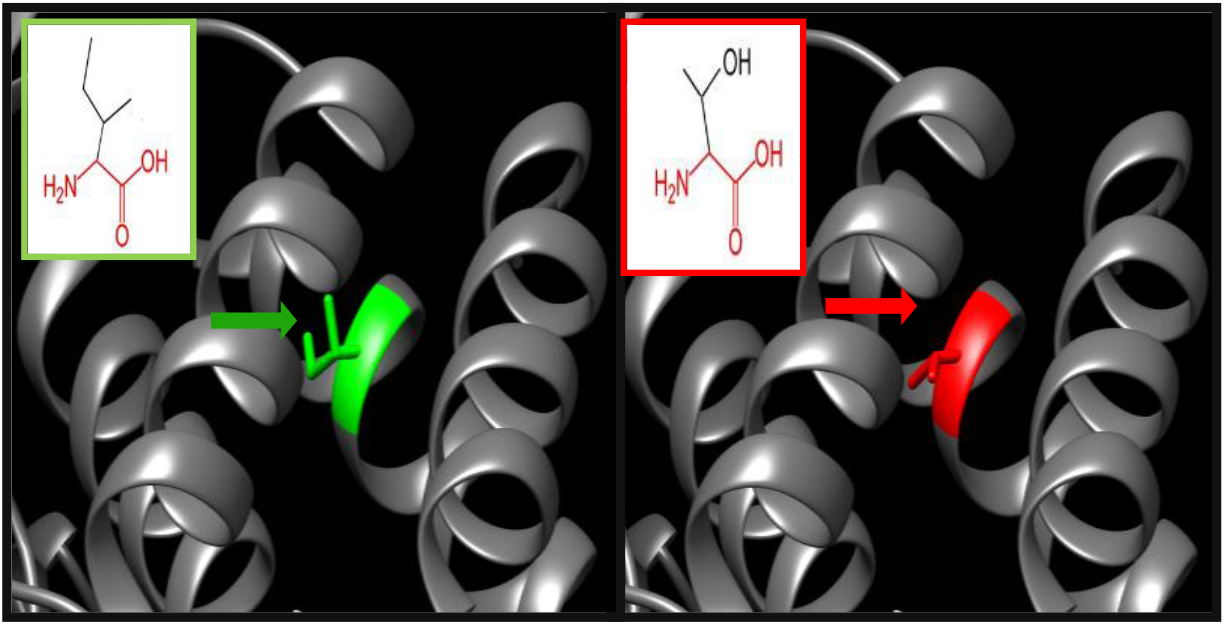
(rs750565047) (I327T) the amino acid Isoleucine changes to Threonine at position 327.

**Figure 22:**
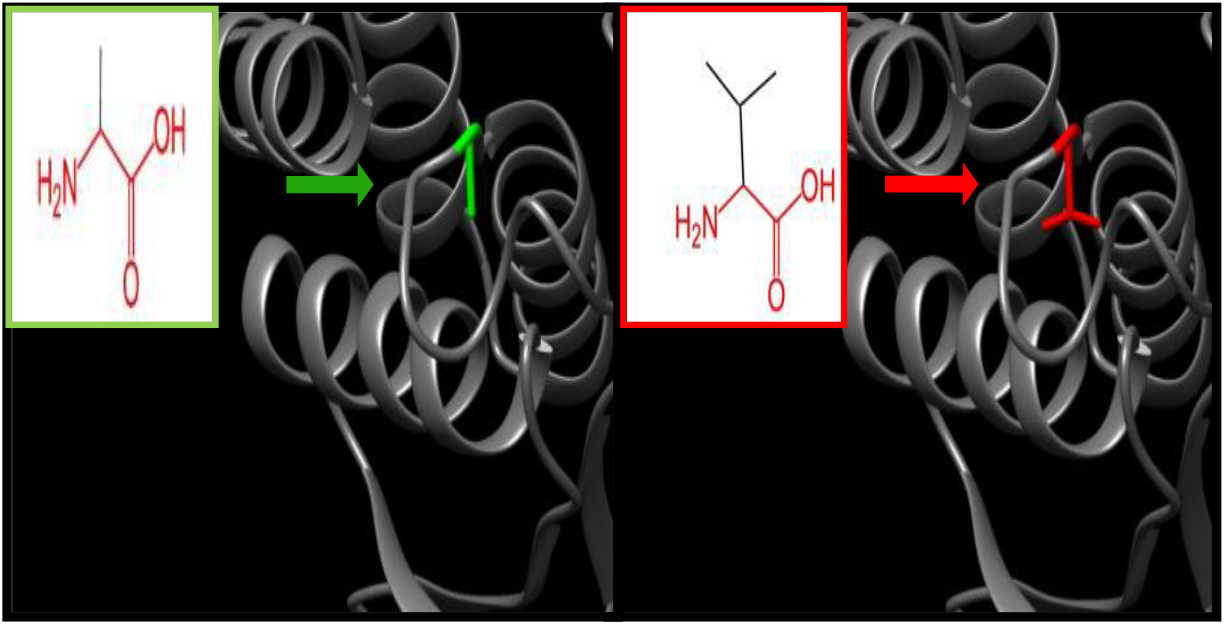
(A302V) the amino acid Alanine changes to Valine at position 302

**Figure 23:**
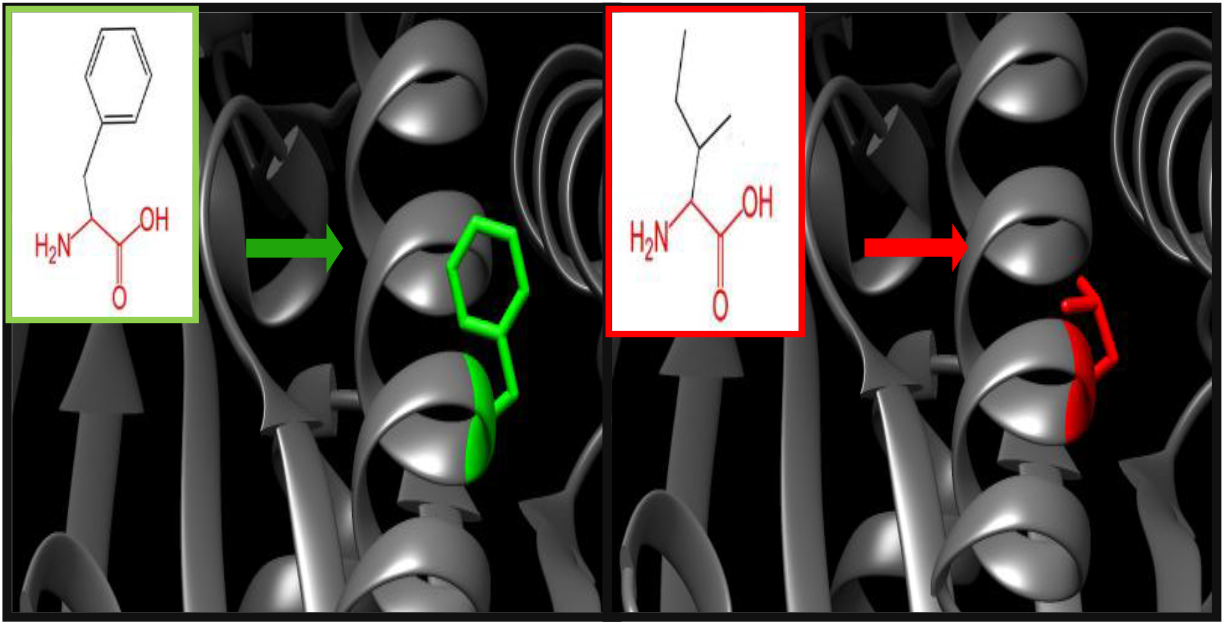
(F184I) the amino acid Phenylalanine changes to Isoleucineat position 184

## 4. Discussion

20 novel mutations have been found to affect the stability and function of *IDH3A* gene using bioinformatics tools. The softwares used were based on different aspects and used different parameters to determine the pathogenecity of the SNPs and to give clues about the effect of the mutation on the molecular level. Multiple softwares were used in this study and the results were compared in order to minimize the errors and confirm the results. (Figure 1)

The total number of the nsSNP −571-in Homo sapiens located in the coding region of the *IDH3A* gene was obtained from the dbSNP/NCBI Database; out of this number 178 nsSNPs (missense mutations) were submitted to SIFT server, Polyphen-2 server and provean server respectively to predict the effect of the SNPs on the function of the protein. SIFT server predicted 91 SNPs to be deleterious, polyphen-2 server predicted 90 SNPs to be deleterious (68 probably damaging and 22 possibly damaging) while Provean server predicted 100 deleterious SNPs.

The triple positive results-which are the ones affecting the function of the protein-were found to be 24 SNPs (see table 1); they were submitted to PHD-SNP, SNP&GO and P-MUT for further analysis. 20 deleterious SNPs were predicted by PHD-SNP and SNP&GO while P-MUT predicted 23 deleterious SNPs, so the triple positive result −20 SNPs-(see table 2) was submitted to I-Mutant and MUpro to investigate their effect on the stability.

The 20 SNPs were all found to be decreasing the stability of the protein except for 2 SNPs (F184L and A302V) which were predicted by I-Mutant to increase the stability of the protein (see table 3).

The sequence of *IDH3A* was aligned against 6 members of its homologues using Bioedit software, and all the SNPs were found to be in a conserved region agreeing with the result obtained from Project Hope which revealed that the SNPs are all found in a domain. This suggests that the SNPs will have an effect on the structure and function of the protein. As an example, P304H mutation was found in a conserved region (see figure 3). Project HOPE uncovered that the mutant (Histidine) is bigger than the wild-type (Proline) and the wild-type was buried in the core of the protein so the mutant will probably not fit. In addition to that, there is a difference in the hydrophobicity between the two and the mutation will cause loss of hydrophobic interactions in the core of the protein. This mutation was confirmed pathogenic in our study and that comes with agreement to another study that confirmed the association of P304H mutation with sever encephalopathy in infancy. (28)

GeneMANIA revealed that *IDH3A* interacts with so many genes and functions in the cellular respiration and energy derivation by oxidation of organic compounds in the tricaboxylic acid cycle, and abnormalities in this function was found to play a key role in the pathogenesis of Bipolar Disorder, therefore *IDH3A* could potentially be a novel therapeutic target for bipolar disorder. (29) The genes co-expressed with, share similar protein domain, or participate to achieve similar function were illustrated by GeneMANIA and shown in figure (2) Tables (4 & 5). UCSF Chimera was used to visualize and analysis of molecular structures. (Figures 4-23)

Also aberrant expression of *IDH3A* promotes malignant tumor growth so could be used as a novel target for the diagnosis and treatment of cancers (30, 31).

Our study identified 20 deleterious SNPs that affect the stability and the function of the protein. All these SNPs were retrieved from the NCBI as untested and this study confirmed them damaging. Our study is the first in silico analysis of *IDH3A* gene. It was based on functional analysis while all previous studies (25, 28) were based on exome sequencing. The 20 deleterious SNPs might be considered as a potential novel target in the diagnosis of Retinitis Pigmentosa and associated diseased.

## 5. Conclusion

In this study the effect of the SNPs of *IDH3A* gene was thoroughly investigated through different bioinformatics prediction softwares. 20 novel mutations were found to have a damaging impact on the structure and function of the protein and may thus be used as diagnostic markers for Retinitis Pigmentosa and could help in the overall understanding of the disease.

## Acknowledgement

The authors wish to acknowledge the enthusiastic cooperation of Africa City of Technology-Sudan.

